# AR suppression rewires a SOX4 regulatory program to promote enzalutamide resistance in prostate cancer

**DOI:** 10.64898/2026.04.27.721035

**Authors:** Sini Hakkola, Anni Perämäki, Antti Kiviaho, Alfonso Urbanucci, Matti Nykter, Mikael Marttinen

## Abstract

Treatment resistance in prostate cancer remains a major clinical challenge. While single-cell transcriptomics enables mapping of cellular composition and resistance-associated states, the gene regulatory programs driving these state transitions remain poorly defined. Here, we integrate single-cell chromatin accessibility and transcriptome sequencing data from an *in vitro* model to resolve the regulatory program architecture underlying anti-androgen resistance. We identify coordinated remodeling of androgen signaling- and stress response-associated regulatory programs, together with a pronounced gain of a SOX4-driven regulatory program in resistant cells. Analysis of clinical datasets reveals that SOX4 activity is both context-dependent and lineage-specific. In primary tumors, SOX4 *cis*-regulatory activity is elevated in PTEN-loss tumors, consistent with a PTEN–PI3K/AKT/mTOR–SOX4 axis that primes AR-independent epithelial lineages for subsequent activation of a more extensive SOX4 *trans*-regulatory program. This program is further amplified following AR-targeted therapy, indicating therapy-induced reinforcement of SOX4-linked regulatory adaptation. TF footprinting additionally implicates Nuclear Factor I family transcription factors as upstream regulators of SOX4, supporting a role in promoting SOX4 activation following AR inhibition. Together, these findings demonstrate that AR inhibition reshapes the gene regulatory landscape to promote engagement of the SOX4 program, facilitating prostate cancer progression under anti-androgen therapy.

## Introduction

Prostate cancer (PCa) cells rely on androgen receptor (AR) signaling, a lineage-defining transcription factor (TF), and primary target in first-line therapies for PCa patients, aiming to reduce circulating androgen hormone levels and ultimately diminish AR activity to castrate levels. Eventually, PCa patients progress to castration-resistant prostate cancer (CRPC) which can be resistant to both androgen deprivation therapy (ADT) and direct therapeutic targeting of AR through AR signaling inhibitors (ARSIs), such as enzalutamide (Enza). While recurrent genetic alterations, including AR amplification or mutation and loss of key tumor suppressors such as RB1, TP53 and PTEN^1,2^ contribute to disease progression, they do not fully explain the phenotypic and transcriptional heterogeneity observed in resistant tumors^3–8^. Resistance to ARSIs or ADT can evolve dynamically under treatment pressure giving rise to diverse cellular states that either restore AR signaling or transition towards alternative lineage programs, such as neuroendocrine (NE) or other AR-independent states^3,7^.

Extensive single-cell and spatial transcriptomic studies have mapped the cellular composition of PCa, its tumor microenvironment, and evolutionary trajectories associated with disease progression^6,9–16^. However, a mechanistic understanding of the TF regulatory processes that drive these state transitions remains limited. Epigenetic remodelling provides a plausible mechanism by which tumor cells rewire transcriptional programs, alter lineage identity, and adapt to sustained therapeutic stress independently of new genetic alterations^17^. Relatively few studies have systematically interrogated the gene regulatory programs of prostate tumors by jointly modeling chromatin accessibility and transcriptional outputs. Bulk multi-omic analyses of advanced PCa have inferred regulatory networks linking chromatin accessibility to gene expression and identified subtype-specific TF motif activity associated with progression and resistance through epigenetic remodeling^7,18^. However, as tumor cells strive in their microenvironment, this complexity limits the ability to disentangle cancer cell-intrinsic regulatory programs from microenvironmental influences. Prior work examining such tumors at single-cell level has identified treatment-persistent subpopulations with distinct chromatin and transcriptional states^19^, and experimental models have captured chromatin and transcriptional adaptations during the evolution of resistance to ARSIs^20^. Yet, these studies have analyzed cell chromatin accessibility and transcriptional profiles as separate modalities, without resolving the architecture of how TF-driven gene regulatory programs are rewired during cancer cells adaptation to treatment.

Here, we reconstruct PCa cell-intrinsic gene regulatory programs underlying treatment response and resistance to second-generation anti-androgen Enza by integrating single-cell chromatin accessibility and transcriptomic profiling of Enza-exposed and -resistant cells. This approach revealed coordinated and hierarchical processes remodeling the androgen signaling- and stress response-associated regulatory programs, alongside the emergence of SOX4-associated resistance-driving programs. We demonstrate that footprints from these regulatory programs predict response to Enza in a PCa patient cohort and exhibit context dependence across both genetic backgrounds and tumor composition. Finally, we identify a potential mechanistic link between AR suppression and altered activity of its binding partners within the Nuclear Factor I (NFI) TF family, providing a framework to explain the gain of SOX4 regulatory programs.

## Results

### Reconstruction of gene regulatory programs in enzalutamide-exposed and - resistant prostate cancer cells

We reasoned that the development of resistance would involve coordinated changes in TF activity and downstream gene expression programs. To investigate anti-androgen resistance-related TF regulatory reprogramming, we reconstructed gene regulatory programs through integration of unpaired scRNA-seq and scATAC-seq data from LNCaP parental cells and two Enza-resistant derivatives (Res-A and Res-B)^20,21^ **(Fig. 1a, b**). To identify candidate *cis*-regulatory interactions, we linked highly variable genes to nearby ATAC peaks and jointly embedded them to a common low dimensional space (Methods, **Supplementary Fig. 1a**). We discovered 3,916 significant *cis*-regulatory peak-to-gene connections (*q*-val 0.1). By integrating these *cis*-regulatory links with TF–gene co-expression patterns and TF motif support, we inferred a total of 44 TF–target gene entities (TF regulons) (**Supplementary Data 1**, Methods), with downstream analyses focusing on the most prominent regulons (n = 23, Methods). To further capture TF activity dynamics, we assigned an activity score to single cells based on the expression of genes assigned to each regulon, providing a cell-by-regulon representation of the data. Single cells were grouped into metacells (**Fig. 1c**, Methods) to explore the regulon behavior across cell states.

**Figure 1.**
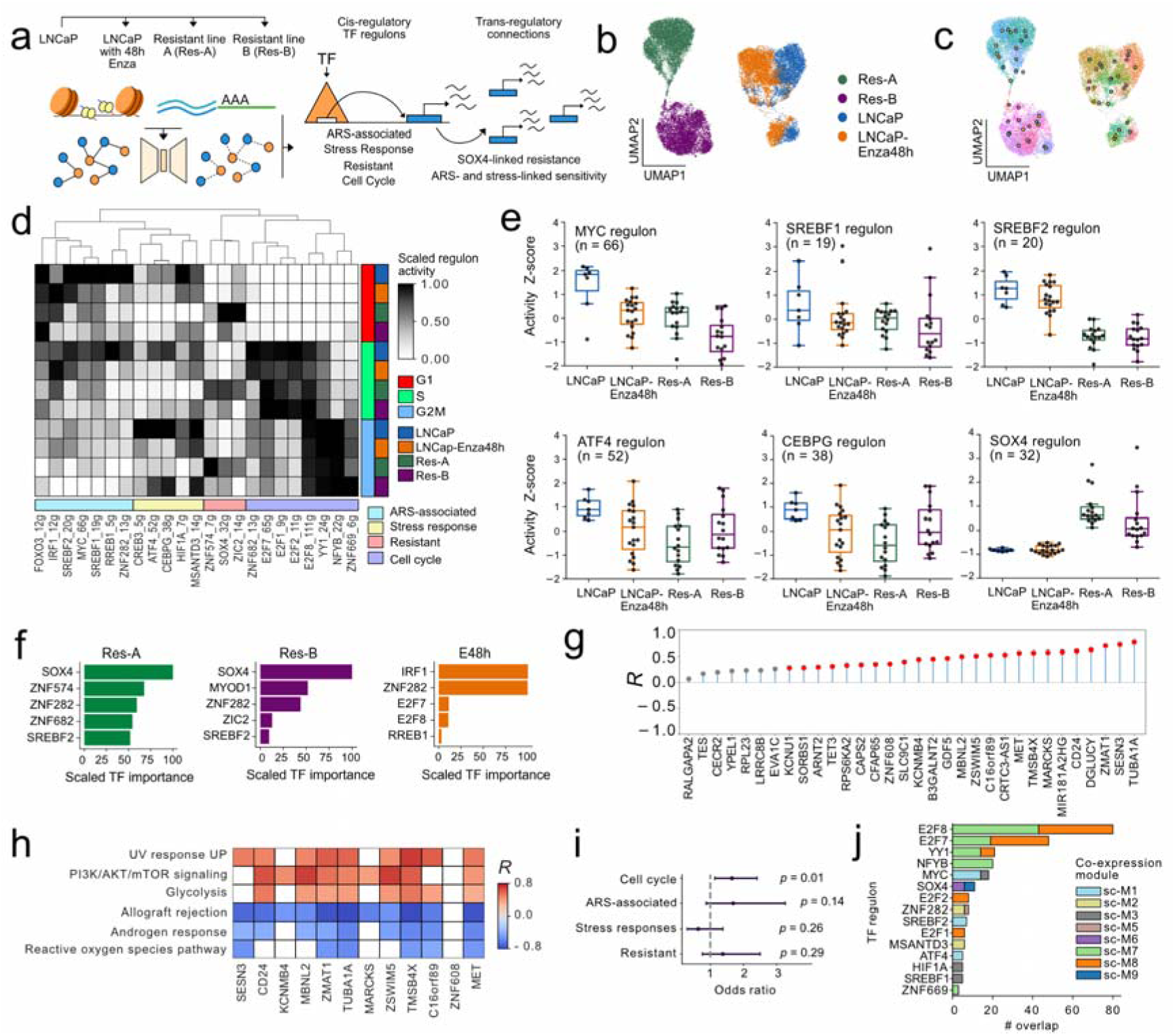
TF regulatory programs identified from single-cell chromatin accessibility and transcriptomic profiling capture enzalutamide resistance-associated reprogramming. **a** Schematic of the analysis workflow for identifying TF regulons and associated *trans*-regulatory connections in Enza resistance. **b–c** UMAP of the ATAC and RNA cells in integrated embedding colored by **b** samples **c** metacell assignment of single cells. Larger points denote metacell centroids computed as the mean UMAP coordinates of member cells. **d** Heatmap of the scaled TF regulon activities across the samples and cell cycle phases, showing TFs expressed in ≥ 2 % of cells with a minimum of five target genes (n = 23). **e** Activity of MYC, SREBF1, SREBF2, ATF4, CEBPG and SOX4 regulons across samples on metacell level. Metacells were grouped based on the dominant sample within each metacell. Values represent the Z-score normalized activity scores within metacells. **f** Top five most important TFs among the full TF regulon landscape (n = 44), ranked by the ability of their regulons to predict differential gene expression in Res-A, Res-B and LNCaP-Enza48h relative to the parental control sample. **g** Lollipop plot of Spearman correlation coefficient between the expression of SOX4 regulon genes and *SOX4*. Red dots denote correlations with *p*_Adj_ < 0.05. **h** Heatmap of Spearman correlation coefficient between SOX4 regulon differentially expressed (log2FC > 1, *p*_Adj_ < 0.05) genes and the MSigDB hallmark gene sets with most significant correlations on the level of metacells. Correlations with *p*_Adj_ ≥ 0.05 are masked. **i** Forest plot of odds ratios (95 % confidence interval) of the enrichment for the differentially upregulated genes in Res-A and Res-B in regulon groups. Enrichment was assessed by comparing the proportion of DEGs within each regulon group between Res-A and Res-B using Fisher’s exact test. Odds ratio > 1 indicates higher relative enrichment in Res-A and < 1 indicates higher enrichment in Res- B. **j** Barplot describing the overlap between TF regulons and co-expression modules with one-sided Fisher’s exact test *p*_Adj_ < 0.05 (alternative hypothesis test ‘greater’ for overrepresentation)

A previously identified ARSI-persistent cell state is marked by the Persist signature^20^ which peaks in S/G2M phases (**Supplementary Fig. 1b, c**) and is associated with poor prognosis through elevated proliferation^22^. This indicates regained proliferative capacity under AR inhibition, increasing the proportion of cells in later cell cycle phases. However, we observed no clear gain or loss of activity for cell cycle-associated regulons (YY1, NFYB, ZNF669, ZNF682 and E2F TFs) between parental and resistant cells **(Fig. 1c**), suggesting that persistent proliferative capacity is instead driven by alternative programs compensating for suppressed AR signaling rather than classical cell cycle controllers.

At the target gene level, we observed largely independent regulons, although some showed partial overlap, suggesting this approach also captures certain coordinated efforts of TFs (**Supplementary Fig. 1d**). For example, ATF4 and CEBPG shared a substantial fraction of their targets, in line with their consistent cooperative role in stress responses^23^.

### Enzalutamide rewires stress signaling and resistance-promoting SOX4-linked gene regulatory programs

Beyond the *Cell Cycle regulons*, we defined three additional regulon groups based on their shared activity patterns, functional enrichments and known biological roles of their TFs. Consistent with AR pathway suppression we identified a group of *Androgen Signaling (ARS)-associated regulons* with reduced activity in Enza-treated parental and resistant cells (**Fig. 1d, e**). These regulons were enriched for androgen and hypoxia signaling and unfolded protein response (UPR) (**Supplementary Fig. 1e**), linking this group to androgen-responsive and stress-adaptive functions. In parallel, we observed repression of *Stress Response regulons*, governed by classical mediators of stress-related signaling, such as ATF4, CEBPG, CREB3and HIFIA1 (**Fig. 1d, e**)^23–25^. Beyond their expected UPR enrichment, these regulons showed substantial overlap with the biological functions defining the ARS-associated regulons (**Supplementary Fig. 1e**), indicating interconnected stress-adaptation and AR-linked programs and multilayered regulatory architecture. Lastly, the resistant stage was marked by increased activity of *Resistant regulons* for ZNF574, ZIC2 and SOX4 (**Fig. 1d, e**).

To determine which TFs within these regulon groups most strongly contribute to the transcriptional differences between parental and resistant cells, we applied a network-based prioritization framework (Methods)^26^. This analysis evaluates how well each TF regulon explains the observed differential gene expression between resistant lines and the parental LNCaP line. Notably, SOX4 emerged as the top regulator in both Res-A and Res-B comparisons, underscoring its potential biological relevance for driving the resistant stage transcriptional adaptations (**Fig. 1 f**), primarily through increased SOX4 expression leading to activation of its target genes (**Fig. 1 g**).

Having mapped the TF regulon-level changes and their associated biological processes, we next proceeded to examine global pathway activity across Enza-treated and resistant states relative to parental cells. Regulon-based metacell clustering revealed two populations: *proliferating* (G2M) and *non-proliferating* populations (G1) (**Supplementary Fig. 1 f**). Because resistant states did not involve major changes in the Cell Cycle regulons, we focused specifically on cells in non-proliferating metacells, to capture alterations preceding S phase without confounding enrichment of S/G2M-related gene sets. Rank-based gene set enrichment analysis (GSEA) on datasets revealed a potential compensatory gain of growth-promoting and metabolic processes such as glycolysis and fatty acid metabolism as well as pro-tumorigenic PI3K/AKT/mTOR (PAM) signaling in resistant cells (**Supplementary Fig. 1 g**). Expression of several SOX4 regulon genes positively correlated with the activity of PAM on the metacell level (**Fig. 1 h**), in line with previous reports connecting SOX4 and PAM activation in PTEN-loss contexts^27,28^. Conversely, expression of SOX4 target genes showed a negative correlation with androgen response (**Fig. 1 h**).

Despite the shared gain in SOX4 regulon activity, we noticed an evident divergence between the resistant lines. While major cell cycle-associated regulatory rewiring did not emerge as a common resistance feature, the two resistant lines differed in their proliferative behavior. Differential expression analysis (resistant line vs. control or Enza-treated parental line) revealed a significant enrichment of Res-A–associated upregulated genes for the Cell Cycle regulons as compared to Res-B (*p* = 0.01, Fisher’s exact test), consistent with the higher proportion of Res-A cells in S/G2M phases (**Fig. 1i, Supplementary Fig 2a**). In contrast, Res-B upregulated genes contributed more strongly to Stress Response regulons. Although ARS-associated and Stress Response regulons were enriched for UPR components (**Fig. 1e**), GSEA did not reveal a significant global enrichment of UPR across samples (**Supplementary Fig. 1f**, showing only the gene sets enriched with *p* < 0.05). Instead, a clear dichotomy in the regulon-associated UPR gene expression was observed, with some genes activated in parental cells and others in Res-B (**Supplementary Fig. 2b**). Our regulon inference identified *XBP1* as part of ATF4 and CEBPG regulons. As a known ATF4 target, *XBP1* promotes cell survival through chaperone-mediated protein folding^29^. *XBP1* was predominantly expressed in Res-B cells (**Supplementary Fig. 1b**), alongside upregulation of an XBP1-dependent chaperon program, indicating rewired protein folding capacity (**Supplementary Fig. 2c**). Together, these findings suggest that resistance involves a shift in UPR composition rather than global increase in UPR activity. Specifically, Res-B cells bias toward a pro-survival, XBP1-driven branch while maintaining a comparable baseline of UPR activity. Thus, the SOX4 regulon emerges as a shared molecular backbone across both resistant cell lines, while stress adaptation and proliferative capacity diverge between Res-A and Res-B.

To further contextualize the ARSI-induced regulatory shift in ARS-associated and Stress Response regulons, we examined the expression and activity of the core TFs underlying these regulons. Most of the TFs showed their highest expression in the parental line (**Supplementary Fig. 2d**). For the stress response-regulators ATF4 and CEPBG, regulon activity strongly correlated with the TF gene expression (R = 0.81, p= 2.43 x 10^-14^ and R = 0.86, p= 3.83 x 10^-18^, Student’s t-test, respectively) (**Supplementary Fig. 2e, f**), consistent with their cooperative, activatory roles in stress-related gene programs^30^. Although highest in parental metacells, subsets of Enza-treated and resistant metacells also retained elevated ATF4 and CEBPG activity (**Fig. 1e, Supplementary Fig. 2e, f**). Together, these observations indicate that ATF4- and CEBPG-mediated stress programs remain broadly engaged across all treatment states, without pronounced shift upon resistance. Instead, the Res-B phenotype reflects a selective downstream reorganization, marked by preferential engagement of the XBP1-driven branch.

By comparison, ARS-associated regulons such as MYC, SREBF1, and SREBF2 exhibited significantly increased motif activity in resistant cell states compared to the parental (MYC: *p* < 0,0001, SREBF1: *p* < 0.001 and SREBF2: *p* < 0.001, Mann-Whitney U test) (**Supplementary Fig. 2g**) despite a concomitant reduction in their transcript levels and decreased expression of several inferred target genes (**Supplementary Fig. 2h-2j**). This discordance suggests a possible compensatory chromatin remodeling: decreasing TF abundance may silence canonical targets, while increases in motif accessibility preserve regulatory potential at essential loci. Because TF regulatory inference captures dominant co-expression patterns, it may not fully resolve targets whose expression is maintained independently of TF transcript levels.

While the TF regulon analysis highlights the *cis*-regulatory changes, it does not fully capture broader transcriptional coordination across resistance states. Finally, to extend these *cis*-regulatory insights gained from the TF regulons, and to capture broader *trans*-regulatory structures, we performed a co-expression analysis (Methods). This analysis identified a co-expression module, sc-M1 overlapping with ARS-associated MYC and SREBF2 regulons as well as the Stress Response regulon ATF4, consistent with cooperative ARS-stress regulatory program (**Fig. 1j and Supplementary Fig. 2k**). In contrast, two co-expression modules (sc-M6 and sc-M9) overlapped significantly with SOX4 regulon (*p*_Adj_ = 1.66 × 10^-5^ and 9.17 × 10^-5^, respectively, one-sided Fisher’s exact test) (**Fig. 1j**), implicating these modules in the SOX4-associated Enza-resistance program. Module sc-M9 was retained for further analyses (**Supplementary Fig. 2l**). To facilitate elevation in clinical cohorts, we derived these findings into two non-overlapping *trans*-regulatory programs: a ARS- and stress-linked sensitivity program (336 genes) and SOX4-linked resistance program (239 genes) (**Supplementary Data 2**, Methods).

### AR/PTEN status and the cell-type context shape SOX4 regulatory activity

To assess the clinical relevance of the ARS- and stress-linked sensitivity program and SOX4-linked resistance program in the context of Enza treatment, we analyzed RNA-seq data from Enza-treated patients classified as responsive or non-responsive based on PSA-levels after 12 weeks^31^. The two *trans*-regulatory programs were predictive of survival in this cohort (**Fig. 2a,b**). Notably, higher ARS-and stress-linked activity was associated with better survival (*p* = 0.005, log-rank test), while higher SOX4-linked resistance activity was predictive of poorer survival (*p* = 0.049). The 13-gene core SOX4 regulon (**Fig. 2c**) was likewise predictive of poorer survival (*p* = 0.018); however, unlike the broader *trans*-regulatory networks (*p* = 0.030 for ARS- and stress-linked resistance and *p* = 0.045 for SOX4-linked resistance, limma moderated t-tests with empirical Bayes variance shrinkage) it did not distinguish PSA-defined responders from non-responders (*p* = 0.603) (**Supplementary Fig. 3a–c**). This indicates that the SOX4 regulon captures additional biology relevant to survival that is not reflected by the PSA status alone.

**Figure 2.**
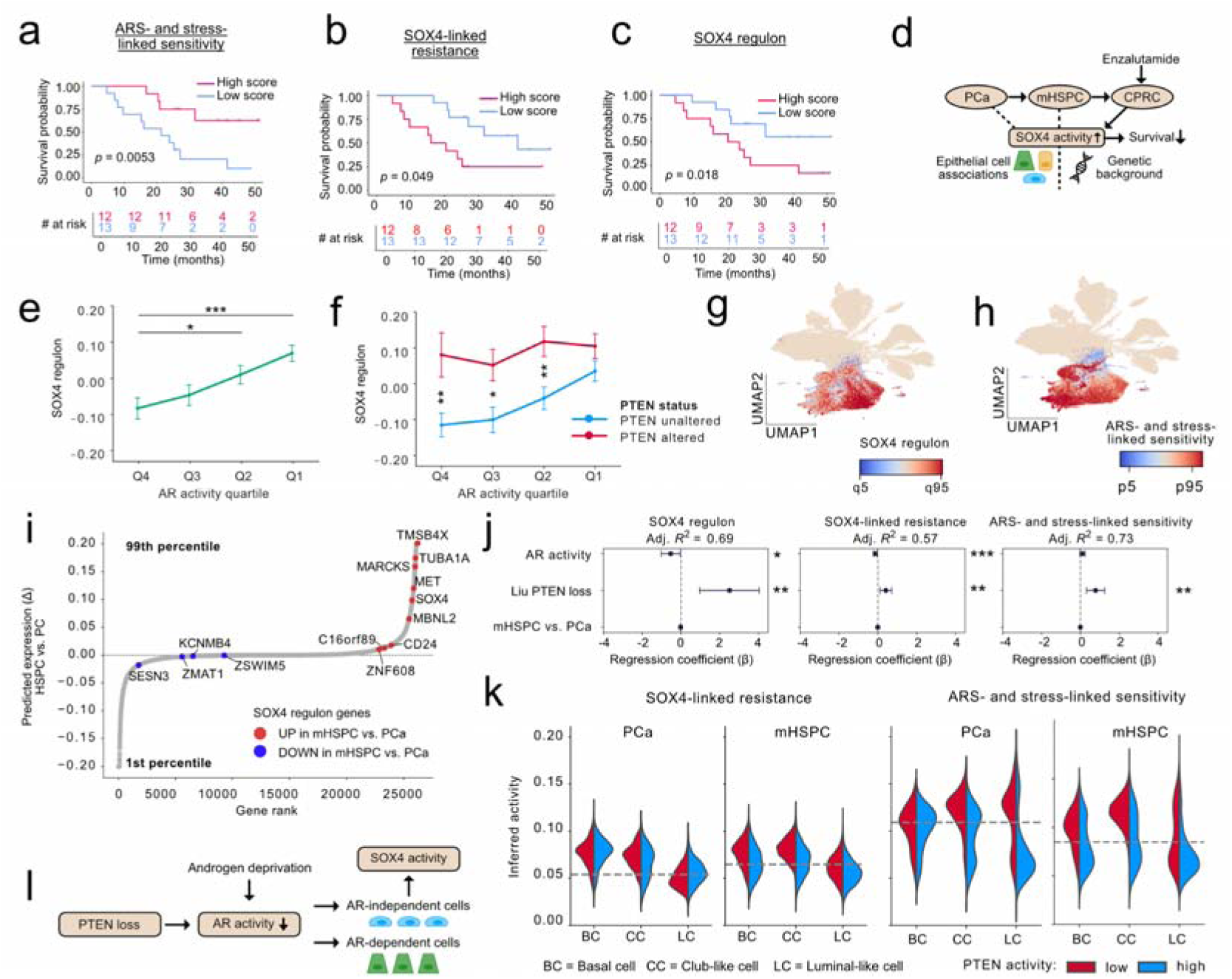
SOX4 program associates with low PTEN activity and AR-independent cell identities. **a–c** Kaplan-Meier survival curves for RNA-seq data from Enza-treated patients (n = 25) with metastatic CRPC stratified into two groups based on median GSVA score (log-rank test *p*-value) for **a** ARS- and stress-linked sensitivity **b** SOX4-associated resistance **c** SOX4 regulon. **d** We aimed to understand whether the SOX4-related programs emerge prior to anti-androgen therapy, and their evolution during PCa progression. **e–f** SOX4 regulon activity in RNA-seq data from TCGA PCa cohort (n = 491) across **f** AR activity quartiles **e** AR activity quartiles stratified by PTEN CNA status (* *p* < 0.05, ** *p* < 0.01, *** *p* < 0.001, Mann-Whitney U test). **g–h** UMAP representation of scRNA-seq data from PCa and mHSPC patients (n = 37) colored by activity of **g** SOX4 regulon activity and **h** ARS/stress-linked sensitivity. **i** Rank plot of predicted expression change (Δ) of SOX4 regulon in mHSPC vs. PCa in scRNA-seq data **j** Ordinary least squares (OLS) regression coefficients for models predicting activity of SOX4 regulon, SOX-linked resistance and ARS- and stress-linked sensitivity using PTEN-loss signature activity, AR activity and tumor type (PCa vs. HSPC). Dots show coefficients and error bars of the 95 % confidence intervals (HC3 robust SE). Scores were pseudo-bulked per patient. Adjusted *R*^2^ is shown for each panel (* *p* < 0.05, ** *p* < 0.01, *** *p* < 0.001, Student’s t-test). **k** SOX4-linked resistance and ARS-and stress-linked sensitivity in PCa and mHSPC epithelial cell sub-lineages stratified by high and low PTEN activity groups based on median PTEN activity score. **l** Our findings indicate that PTEN alterations create a permissive state enabling SOX4 regulatory program activation and expansion upon AR-targeted therapy, particularly in AR-independent epithelial cell lineages.

Because the ARS- and stress-linked program predicted treatment sensitivity whereas the SOX4 regulon and SOX4-linked program activity were associated with resistance and poorer survival, we next sought to define the genetic and cellular contexts in which these programs arise and how they relate to one another. Specifically, we asked whether SOX4-linked activity emerges prior to anti-androgen therapy, and how it evolves during disease progression (**Fig. 2d**). Given the central role of AR signaling PCa progression, we first scored untreated prostate tumors^32^ using a previously defined AR activity signature^32,33^. Consistent with prior reports showing that AR can suppress *SOX4* expression^34^, SOX4 regulon activity increased as AR activity declined (**Fig. 2e**). Prior studies have also implicated SOX4 in PCa tumorigenesis particularly upon PTEN loss and resulting PAM pathway activation. Intriguingly, stratification by PTEN copy number status revealed that PTEN loss permits elevated SOX4 regulon activity even in tumors with high AR activity (**Fig. 2f**). As expected, PAM activity was likewise higher in PTEN-altered tumors (**Supplementary Fig. 3d**). In contrast, ARS- and stress-linked signature was closely coupled to AR signaling, with a PTEN-associated increase observed in tumors within the lowest AR activity quartile (**Supplementary Fig. 3e**).

To further contextualize these relationships, we turned to scRNA-seq data from treatment-naïve PCa and metastatic hormone-sensitive disease (mHSPC) on ADT (**Fig.**□**2g–h**; **Supplementary Fig.**□**3f–g**). Using a cluster-free, multicondition differential expression model, we found that most SOX4 regulon genes were more highly expressed in mHSPC than primary PCa cells (**Fig. 2i)**. Beyond copy number loss and mutations, PTEN protein expression can be silenced through transcriptional and post-translational mechanisms^28,35–37^. The transcriptional effects of PTEN loss have been captured in a gene signature^38^. We applied this signature to classify the PTEN status in both PCa and mHSPC patient samples. Linear regression analysis indicated PTEN loss was a stronger determinant of core SOX4 regulon than AR activity (PTEN: *p* = 0.001, 95% CI [1.04, 4.27]; AR: *p* = 0.041, 95% CI [−0.90, −0.02], Student’s t-test). Both PTEN loss and AR activity positively predicted ARS- and stress-linked sensitivity, with PTEN showing stronger association (*p* = 0.040, 95% CI [0.14, 1.29] and *p* = 0.014, 95% CI [0.01, 0.30], respectively). (**Fig. 3j**). While the SOX4-linked resistance also showed alink to both low AR and high PTEN loss activity (AR: *p* = 0.041, 95% CI [-0.21, -0.05]; PTEN: *p* = 0.001, 95% CI [0.12, 0.68]) this relationship was primarily driven by the core SOX4 regulon activity (**Supplementary Fig. 3h**), consistent with a model in which the PTEN**–**PAM axis primes the PCa cells for activation of broader SOX4 program activation upon AR-targeted therapy. Tumor type itself did not significantly contribute to the model (**Fig. 3j**), indicating that differences between PCa and mHSPC largely reflect a higher prevalence of PTEN-low or AR-low states in mHSPC.

**Figure 3.**
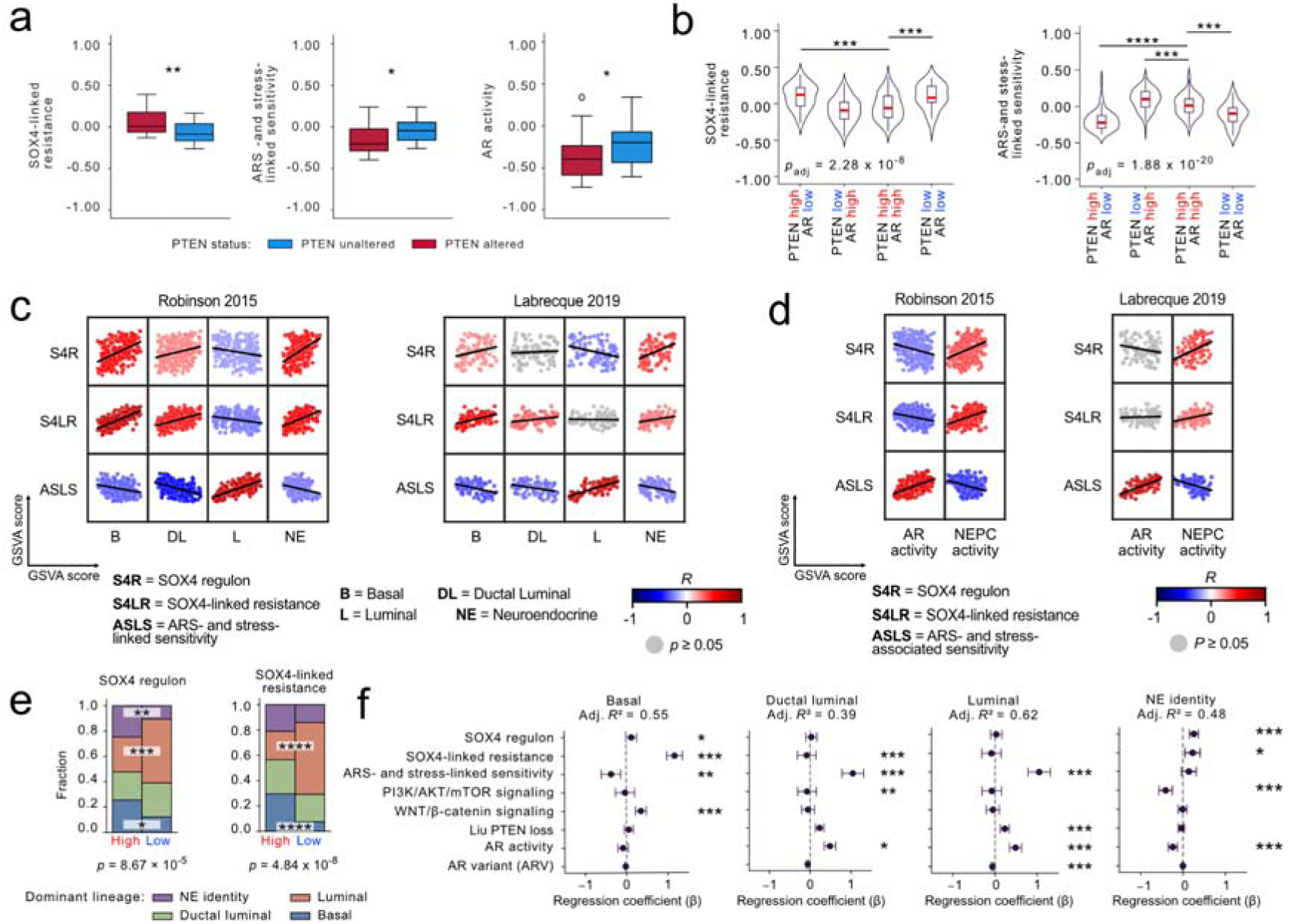
SOX4 regulatory program is associated with AR independency while ARS- and stress-linked program points to AR-dependent states in CRPC. **a** GSVA score of SOX4-linked resistance, SOX4 regulon, ARS- and stress-linked sensitivity and AR activity in tumors from Labrecque 2019 cohort presented as boxplots, stratified by PTEN copy number status, PTEN altered (n = 26) and PTEN unaltered (n = 20), was sourced from Roudier *et al* 2025. Asterisks represent Mann-Whitney U test *p* * < 0.05 and *p* ** < 0.01. **b** GSVA score of SOX4-linked resistance, and ARS- and stress-linked sensitivity in the SU2C cohort (n = 249) stratified by AR and PTEN activity status. The groups were defined based on the Z-scored GSVA scores (AR low/high corresponded to Z ≤ 0/> 0. As the PTEN signature reflects PTEN loss, higher scores indicate reduced PTEN activity, thus z ≤ 0/> 0 corresponds to PTEN high and PTEN low, respectively. The Kruskal-Wallis *p*_adj_ is displayed (*** *p*_adj_ < 0.001, **** *p*_adj_ < 0.0001, Mann-Whitney U-test). **c–d** Correlation scatter plots comparing the activity of the core SOX4 regulon (S4R), SOX4-linked resistance (S4LR) and ARS- and stress-linked sensitive (ASLS) programs and **c** the epithelial cell subtype-derived signature scores and **d** AR and NEPC activity signature scores in the tumors from SU2C and Labrecque 2019 cohorts (n = 98). The correlations with *p*_adj_ ≥ 0.05 are masked with grey. **e** Epithelial lineage composition in SU2C tumors stratified by SOX4 regulon and SOX4-linked resistance activity. Tumors were divided into High and Low groups based on the z-scored GSVA scores. For each tumor, the dominant epithelial lineage was assigned by the highest z-scored lineage signature. Stacked barplots show the proportional distribution of these lineages within each group. The chi-squared test *p*-value is displayed (** *p*_adj_ < 0.01, *** *p*_adj_ < 0.001, **** *p*_adj_ < 0.0001, Fisher’s exact test). **f** OLS regression coefficients for models predicting the activity of epithelial cell subtype-derived signature scores in the SU2C cohort (* *p* < 0.05, ** *p* < 0.01, *** *p* < 0.001, t-test p-value).

### ARS/stress- and SOX4-linked programs converge in AR-independent epithelial states

At the single-cell level, the SOX4-linked resistance activity was highest in AR-independent club-like and basal cells under PTEN-low conditions across both PCa and mHSPC, with higher activity in mHSPC (**Fig.** □**2k, Supplementary Fig.** □**3i**). Consistently, activity of the core SOX4 regulon localized predominantly in club-like and basal cells (**Supplementary Fig. 3j**). In contrast, the ARS-and stress-linked program exhibited weaker lineage specificity in treatment-naïve PCa and instead segregated primarily by PTEN status (**Fig.**□**2k**). Under ADT, mainly targeting AR-dependent luminal-like cells, ARS- and stress-linked activity remained concentrated in PTEN-low basal and club-like states. Interestingly, we observed a strong positive correlation (*R* = 0.58) between the SOX4-linked resistance and the ARS- and stress-linked sensitivity programs within mHSPC basal and club-like cells (**Supplementary Fig. 3k**). Gene set enrichment analysis of the SOX4-high vs. SOX4-low (Q4 vs. Q1) club-like and basal cells revealed signatures related to mitochondrial metabolism, enhanced protein synthesis and stress- and immune related processes (**Supplementary Data 5**). Activation of the ARS- and stress-linked sensitivity program in these AR-independent epithelial states therefore points to stress response under therapy rather than canonical AR-driven biology. Collectively, these findings support the presence of a metabolically active, stress-adapted SOX4-high subpopulation of basal and club-like cells, consistent with our earlier work linking prostate club cells to immunosuppression and demonstrating that basal and club cells persist under ADT^15^.

Together, these findings indicate that PTEN loss creates a permissive regulatory context for further activation of SOX4-driven transregulatory program upon AR-targeted therapy, particularly in AR-independent epithelial lineages (**Fig. 2l**), whereas the ARS- and stress-liked sensitivity program showed weaker lineage specificity, and instead was more dependent on AR and PTEN status.

### SOX4 regulatory activity aligns with neuroendocrine lineages in CRPC

Our findings support a model in which PTEN loss establishes a permissive regulatory state, allowing the SOX4-linked resistance program to emerge and further expand under ADT. Given the association of SOX4 activity with poorer survival in Enza-treated disease, we next sought to determine how these relationships are further modified in CRPC tumors, the clinical context in which anti-androgens are typically used. Because single-cell datasets of CRPC remain limited, we asked whether our findings were recapitulated in bulk-tumor transcriptomes, where the most clinical data exists.

Using RNA-seq data from CRPC tumors, we first assessed whether associations with PTEN-low and AR-low states persisted. In the Labrecque *et al*, 2019 cohort, using the PTEN alteration status inferred by Roudier *et al*, 2025, both the core SOX4 regulon and SOX4-linked resistance remained elevated in PTEN-altered tumors (*p* = 0.03 and *p* = 0.009, respectively, Mann-Whitney U test) (**Fig. 3a**, **Supplementary Fig. 4a, b**). These tumors also exhibited reduced AR activity (*p* = 0.04), whereas PAM signaling activity did not significantly differ between groups (*p* = 0.991). This likely reflects the frequency of PTEN loss characteristics in CRPC, rather than a persistent coupling between SOX4 regulatory program and PTEN**–**PAM at this disease stage and instead points to a stronger association to AR-low status. About 40-50% of CRPCs are estimated to harbor loss of PTEN, compared with 15-20% in primary PCa^32,39^. In line with a more pronounced effect of AR activity status, the ARS- and stress-linked sensitive program tracked with AR activity rather than PTEN status (**Fig. 3a**). Consistently, in the SU2C cohort the SOX4-linked resistance program stratified more strongly by AR activity than by PTEN status (**Fig. 3b**). SOX4-linked resistance was preferentially activated in AR-low tumors irrespective of PTEN status, whereas the ARS- and stress-linked sensitivity program remained characteristic of AR-high tumors. Together, these findings indicate that although PTEN loss helps establish the SOX4-permissive state in earlier disease, in advanced CRPC the activity of the SOX4-linked program is predominantly shaped by AR-low states, while the ARS- and stress-linked sensitivity program is strongly coupled to high AR activity.

Single-cell data from patients with primary treatment-naïve PCa and mHSPC on ADT indicated that SOX4 activity spans multiple epithelial cell states, particularly basal and club-like lineages. Consistent with this, in the SUC2 RNA-seq cohort, the core SOX4 regulon and the SOX4-linked resistance program correlated positively with ductal-luminal (club-enriched) signature ^40^, and, intriguingly, also with NE-like cell signature (**Fig. 3c**). Similar trends were observed in the Labrecque 2019 cohort where both gene sets correlated positively with the NE-like and basal signatures, and SOX4-linked resistance additionally showed a positive association with the ductal luminal signature (**Fig. 3c**). Conversely, both gene sets exhibited negative or non-significant correlation with luminal-like signatures in both datasets. As expected, both the core SOX4 regulon and the SOX4-linked resistance program also correlated positively with neuroendocrine PCa (NEPC) activity^41^ and negatively with AR activity (**Fig. 3d**). In contrast to the SOX4-associated programs, the ARS and stress-associated sensitivity program was positively correlated with luminal-like signatures and high AR activity and negatively associated with AR-independent cell states and NEPC activity (**Fig. 3c, d**. To complement these continuous correlations, categorical lineage enrichment analysis showed a marked overrepresentation of basal identity and depletion of luminal identity among the tumors with high SOX4-linked resistance program activity, while tumors with high SOX4 regulon activity were additionally enriched for NE-like identity (**Fig. 3e**). These activity patterns were also evident in scRNA-seq data sourced from six CRPC tumors (**Supplementary Fig. 4c–h**). While the SOX4-linked activity was predominantly associated with non-luminal lineages (**Supplementary Fig. 4e–h**), a subset of luminal-like cells also exhibited high levels of SOX4 regulon and SOX4-linked resistance activity (**Supplementary Fig. 4g–i**). Notably, SOX4-linked resistance activity showed positive correlation with AR signaling within luminal-like cells (*R* = 0.25) and GSEA analysis revealed a significant enrichment of the androgen response pathway in the SOX4-high luminal-like subset (NES = 2.35, FDR *q*-val < 0.05). These findings suggest that AR-mediated suppression may not fully eliminate SOX4-linked activity, which can persist and vary within AR-active cells in prostate-derived CRPC tumors.

Although bulk- and single-cell-level correlation patterns revealed clear lineage associations, they did not establish whether the regulatory programs contribute independently of canonical oncogenic pathways. To address this, we performed a multivariate linear regression modeling epithelial lineage score (basal, ductal luminal, luminal, NE-like) using our regulatory programs together with key pathways implicated in both PCa progression, and activity of our regulatory programs, including the PTEN/PAM axis, WNT/β-catenin signaling^42,43^, AR and AR variant activity (**Fig. 3f**). In this model, the SOX4-linked resistance program significantly predicted the ductal luminal and basal states (*p* =1.51 × 10^-10^, 95% CI [0.543, 1.021] and *p* = 2.42 × 10^-29^, 95 % CI [0.940, 1.337], respectively, Student’s t-test), and was also predictive of NE-like state (*p* = 0.01, 95% CI [0.054, 0.417]). The core SOX4 regulon, also predictive of basal state (*p* = 0.03, 95% CI [0.016, 0.281]), was predominantly associated with NE-like activity (*p* = 1.943427 × 10^-6^, 95% CI [0.152, 0.365]), highlighting basal and NE states as points of convergence for the SOX4-driven programs. The ARS- and stress-associated sensitivity program was predictive of luminal identity (*p* = 1.27 × 10^-14^, 95% CI [0.778, 1.308]) and, notably, accounted for more variation than AR or ARV activity (*p* = 1.23 × 10^-11^, 95% CI [0.346, 0.627] and *p* = 1.88 × 10^-7^, 95 % CI [-0.087, -0.039], respectively).

These results indicate that SOX4 and ARS- and stress programs represent a distinct transcriptional axis of epithelial state variation, contributing meaningfully alongside canonical pathways associated with PCa biology. Indeed, low AR transcriptional activity in CRPC tumors has been implicated in the development of Enza resistance^31^. In light of our results, the ARS and stress-associated sensitive connections marks higher AR-dependence and may therefore exhibit comparatively more favorable outcomes. In contrast, the SOX4-high program linked to AR-independent and NE-like states appears to reflect a therapy-persistent trajectory associated with poorer prognosis.

### Identification of NFI factors as putative regulators of SOX4 upon AR inhibition

The core SOX4 regulon and the *trans*-regulatory SOX4-linked resistance program were mainly linked to AR-independent cell states in CRPC. Therefore, the induction of the SOX4 regulatory program upon AR-targeted therapy prompted us to investigate which upstream regulators may activate *SOX4* under therapeutic pressure (**Fig. 4a**).

**Figure 4.**
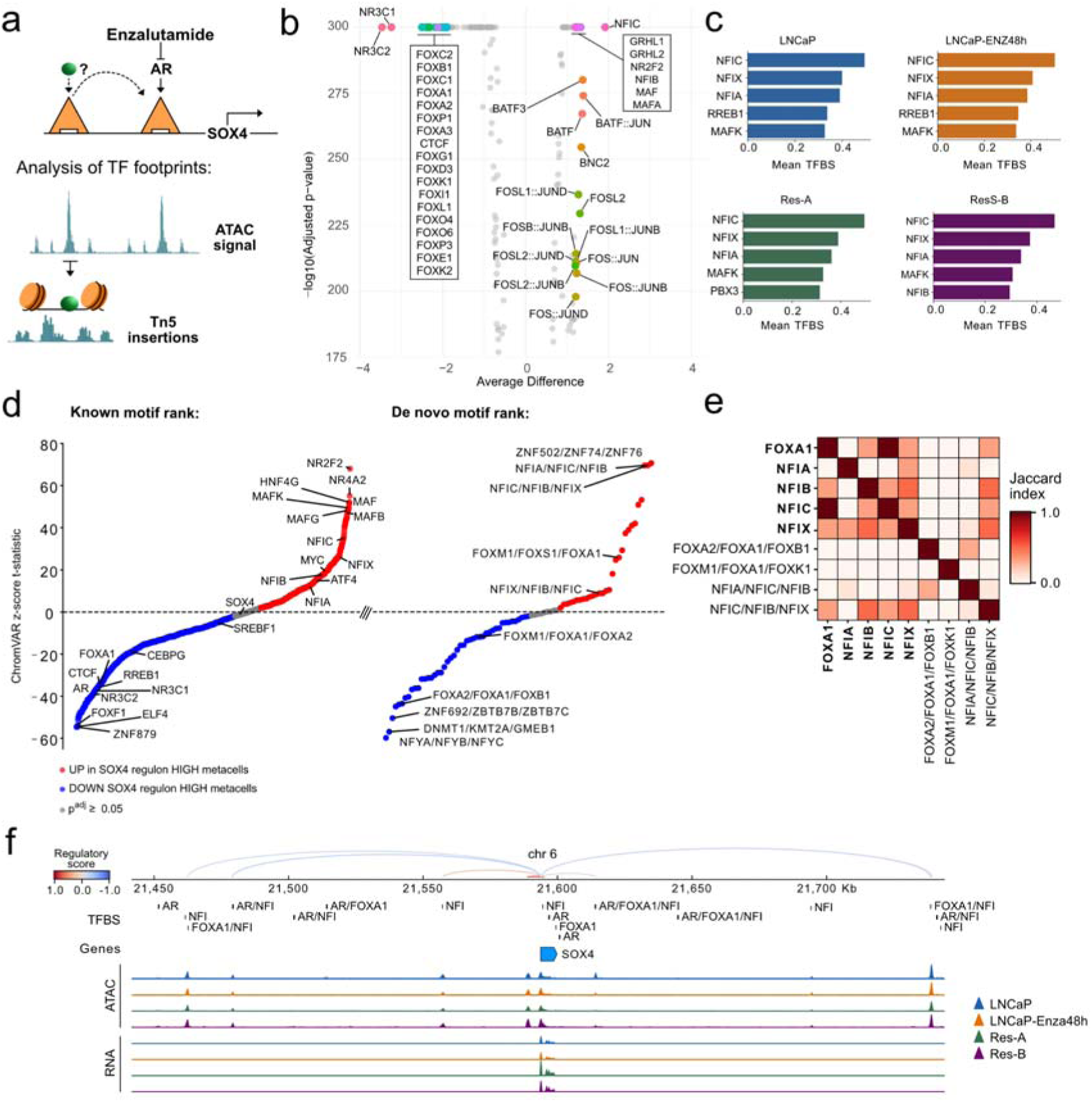
NFI factors as putative regulators of SOX4. **a** We applied TF footprinting analysis on the scATAC-seq data layer from our Enza-resistant *in vitro* model to investigate putative upstream regulators of SOX4 under AR inhibition. **b** Volcano plot of differential motif activity between resistant (RES-A, RES-B) vs. parental LNCaP. cells. Average difference denotes the average motif activity z-score between groups. For visualization purposes, only top 10 down and up regulated motifs based on average difference, with log10(Bonferroni-adjusted *p*-value) ≥ 170. **c** TFs with the highest mean TF binding score (TFBS) in the *SOX4* regulatory region across treatments. TFBS denotes the inferred probability of TF binding at the region. **d** Rank plot of ChromVAR z-score *t*-statistic of the known and *de novo* motifs in metacells with high vs. low SOX4 regulon activity. Each point represents a motif, with red indicating higher accessibility in SOX4 regulon high (Q4) metacells and blue indicating lower activity relative to metacells with low SOX4 regulon activity (Q1). Grey dots indicate *p*_adj_ ≥ 0.05. **e** Jaccard index heatmap describing the co-occupancy of NFI factor and FOXA1 motifs and matching *de novo* motif patterns at peaks in the SOX4 regulatory region. The known motifs are shown in bold. The *de novo* patterns are named after the three most significantly similar known motifs, ranked by FDR *q*-value. **f** Top: The peak–gene links colored by the *cis*-regulatory score between the ATAC peaks and the *SOX4* gene, with the known and *de novo* NFI and FOXA1 TF binding sites (TFBS) and AR motif matches. Co-occurring TFBS and motif occurrences within the same 100bp genomic window are represented as combined labels. Bottom: scATAC-seq and scRNA-seq tracks split by dataset.

Differential motif activity analysis between resistant and parental cells revealed marked loss of Glucocorticoid receptor (GR, NR3C1) and mineralocorticoid receptor (MR, NR3C2) motif activity in resistant cells, suggesting that MR- or GR-mediated hijacking of AR program is unlikely to represent the primary mechanism of resistance in this context (**Fig. 4b**), consistent with the suppression of androgen signaling in the resistant cells (**Supplementary Fig. 1h**).

Previous work linked Enza resistance to increased chromatin accessibility^20,44^. In line with this, we observed a reduction in CTCF motif activity in the resistant cells (**Fig. 4b**). Given CTCFs role as a primary insulator, this could reflect resistance-associated alterations in chromatin architecture^45^. Additionally, several FOX TFs were found among the most downregulated motifs, including FOXA1, an AR co-regulator and a modulator of its transcriptional program^46,47^ and FOXA2, recently found to cooperate with FOXA1 in shaping the AR transcriptome^48^. In contrast, motifs for the NFI family members NFIC and NFIB were strongly upregulated in resistant cells (**Fig. 4b**). Intriguingly, this family has been found to interact with FOXA1 to regulate prostate-specific gene expression via AR and transcriptional reprogramming in Enza-resistant contexts^49,50^. As AR has been reported to repress SOX4^34^, our findings raised the possibility that the NFI factors may activate SOX4 when AR and FOXA1 activity is lost.

To further investigate this, we applied a deep learning-based TF footprinting to chromatin accessibility data from our *in vitro* model. Strikingly, NFI factors showed highest binding probability at the *SOX4* regulatory region in both parental and resistance cells (**Fig. 4c**), suggesting stable chromatin occupancy even under AR inhibition. Consistent with this, *de novo* motif discovery identified multiple NFI and FOX patterns (**Supplementary Data 3**). Both de novo and known NFI motifs were enriched in SOX4 regulon-high metacells (**Fig. 4d**). In contrast, SOX4 motif activity itself was not significantly altered, supporting the notion that increased *SOX4* regulatory program is more closely associated with increased *SOX4* expression than with altered SOX4 motif accessibility. While the *de novo* motif analysis did not reveal composite NFI–FOXA1 motifs, we instead found out that the NFI and FOXA1 motifs co-occupy the same peaks within the *SOX4* regulatory region (**Fig. 4e**), indicating potential co-operative regulation.

Expression level of the *NFI* factor genes appeared relatively stable across the cell model, except for *NFIB*, which showed lower signal on the UMAP in the resistant cells (**Supplementary Fig. 5b**). In clinical single-cell datasets, the expression was concentrated in club-like and basal cells in PCa and mHSPC, where the expression of *SOX4* and its targets was also highest (**Supplementary Fig. 3j**), but also in endothelial cells and fibroblasts (**Supplementary Fig. 5c**). In CRPC scRNA-seq data, the expression of *NFIC* was also elevated in the NE-like and luminal cells (**Supplementary Fig. 5d**). TF footprinting did not reveal AR binding activity in the *SOX4* regulatory region. To assess whether the analysis might miss these events due to the presence of low-affinity AR half-site motifs, we examined whether AR motif matches could be identified within peaks located in the *SOX4* regulatory region. Motif enrichment analysis with Homer^51^ did not reveal enrichment for any TF motif, likely due to the low number of peaks (n = 24). However, a direct motif matching identified several AR motifs within the *SOX4* regulatory region (**Supplementary Fig. 5a**), including those within close proximity (100b) of NFI and/or FOXA1 binding sites (**Fig. 4f**), consistent with potential regulatory co-localization.

Taken together, these findings suggest that AR-targeted therapy reshapes the chromatin landscape to become permissive for alternative TFs such as NFI factors, enabling activation of *SOX4*. In resistant conditions, decreased accessibility at AR and FOXA1 motifs, coupled with increased accessibility at NFI motifs, reflects a shift from canonical AR-driven regulatory programs towards alternative TF networks. This reprogramming may represent a step in establishing AR-independent therapeutic resistance.

## Discussion

In this study, we reconstructed gene regulatory programs in a controlled *in vitro* model of ARSI response and resistance, enabling interrogation of potential epigenetic drivers independently of major genomic confounders^21^. Integration of single-cell chromatin accessibility and transcriptomic data revealed coordinated repression of androgen signaling (ARS)- and stress-linked regulons upon ARSI treatment, a state largely sustained in resistant cells. Resistance was not merely characterized by survival in ARSI-induced repression, but by the emergence of distinct regulatory programs, most prominently the gain of a SOX4-centered program associated with survival-related pathways.

Despite reduced TF expression and downregulation of many inferred target genes, ARS-associated regulators such as MYC and SREBF family members exhibited increased chromatin accessibility at their binding sites in resistant cells. This inverse relationship between TF expression and motif accessibility suggests a compensatory epigenetic mechanism, potentially preserving regulatory potential at essential loci. While such effects are not fully captured by co-association–based network inference approaches, sustained activation of established MYC target pathways supports the notion that chromatin remodeling may buffer transcriptional repression and maintain core proliferative or metabolic programs during resistance.

Most notably, we identified a gain of SOX4 regulon as a defining feature of the resistant state. SOX4 activation was accompanied by increased PAM pathway activity, a well-established signaling axis implicated in therapy resistance across cancer types^52^. The SOX4 program displayed context-dependent associations with AR and PTEN status. In earlier disease stages SOX4 activity was enriched in basal and club-like epithelial populations and associated with PTEN loss, suggesting a PTEN–PAM–SOX4 axis that may prime tumors for development of resistance. Several inferred SOX4 targets, including *MARCKS*, *CD24*, *TUBA1A*, *TMSB4X*, and *MET*, are known regulators of cytoskeleton dynamics and cell motility^53–57^, suggesting a SOX4-driven coordination of cytoskeletal remodeling accompanying the cell state transition.

Despite previous AR-targeted therapy, most CRPCs exhibit active AR signaling enabled by aberrations of the AR gene^58^. Alternatively, tumors can develop to AR-independent or neuroendocrine states^3,8,59^. In advanced CRPC, activation of the SOX4 regulatory program appeared more strongly linked to reduced AR activity and was observed within neuroendocrine-like and other lineage-plastic states. These findings indicate that SOX4 operates within distinct biological contexts during disease progression, shifting from an PTEN-associated program in earlier stages to an AR-deprivation–associated program in advanced disease. Importantly, both the ARS- and stress-linked and SOX4-linked regulatory programs were predictive of treatment response in patients, underscoring their clinical relevance. While the ARS- and stress-linked network marked AR-dependent states and associated with comparatively favorable outcomes, the SOX4 program aligned with AR-independent trajectories and poorer prognosis, consistent with a more plastic and therapy-persistent phenotype.

Finally, our analysis suggests a potential mechanistic link between AR inhibition and SOX4 activation. Ligand-bound AR has been reported to suppress SOX4 expression, with upregulation observed following prolonged androgen deprivation and in CRPC^34,60^. Our data extends this model by indicating that AR-targeted therapy may establish a permissive chromatin landscape enabling alternative TFs, including members of the NFI family, to activate SOX4. Disruption of the AR–FOXA1–NFI regulatory balance could thereby alter TF binding dynamics and facilitate the emergence of resistance-associated regulatory programs. Importantly, SOX4 sits at the intersection of major signaling axes implicated in PCa progression: AR signaling, PAM upon PTEN loss^27,28^, and Wnt/β-catenin^42,43^. Through its ability to respond and modulate these pathways and promote NE-like phenotype^61^, SOX4 may function as a central molecular hub enabling the tumor cells to shift away from AR dependence and adopt alternative therapy resistant phenotypes. Given the metastatic origin of the *in vitro* model and the predominance of metastatic samples among the CRPC validation cohorts, the identified regulatory programs primarily reflect the biology of advanced metastatic CRPC. Elucidating how the SOX4 regulatory program is shaped during tumor evolution and dissemination will further clarify its role in PCa progression and therapy resistance.

Collectively, this work uncovers tumor cell-intrinsic non-mutational regulatory reprogramming during the emergence of PCa treatment resistance. By disentangling genetic and epigenetic contributions in a controlled system and validating findings in human datasets, we provide evidence that epigenetic remodeling plays a central role in shaping adaptive regulatory networks during AR-targeted therapy. These findings support a model in which AR inhibition not only suppresses androgen signaling but actively rewires the hierarchy of gene regulatory programs, enabling alternative survival pathways that drive therapy resistance.

## Methods

### Preprocessing and quality control of the single-cell sequencing data

The FASTQs of scRNA-seq and scATAC-seq data were processed into feature counts using Cell Ranger v.7.1.0 and Cell Ranger ATAC v. 2.1.0, respectively. Peak calling for ATAC-seq data was carried out with *macs3* v.3.0.0b1 with following parameters: *macs3 callpeak -f BED -g hs -q 0.05 --nomodel --nolambda --extsize 200 --shift -100 --call-summits -B -SPMR*. A consensus peak set was generated using an iterative removal process described by Corces MR et al 2018^62^ from peak summits extended to fixed width of 501 bp. Peaks extending beyond the chromosome ends, falling on ENCODE blacklisted genomic regions, observed in only one sample and < 5 cells, and those with a score per million values ≤ 5 were excluded.

Further processing of the data was done in R v. 4.3. Quality control of the scRNA-seq data was conducted with Seurat v.4.3.0.1^63^, excluding genes expressed in ≤ 15 cells and cells with ≤ 1500 or ≥ 8000 features or 60000 UMI counts and ≥ 20 % of mitochondrial reads. For doublet removal, dimension reduction and clustering was performed. Doublets were removed using scDblFinder v. 1.19.6^64^. Signac v. 1.14.0^65^ was used for the quality control of the scATAC-seq data, keeping fragments in ≥ 10 cells and cells with > 1500 and < 30000 peak counts, ≥ 2000 fragments and > 20 % reads in peaks with nucleosome signal < 4 and TSS enrichment > 3.

### Inference of *cis*-reghulatory peak**–**gene connections and TF regulons

The single-cell data was then loaded to Python 3.9 for further analyses. Raw matrix normalization and dimensionality reduction of data modalities was done with scanpy v. 1.9.1^66^. For the RNA layer, 4,000 highly variable features (HVFs) were selected for principal component analysis (PCA), with 50 principal components (PCs) retained. Each cell was normalized by total counts over all genes, then logarithmized and scaled, by centering each gene to have a mean of 0 and scaled to unit variance. Latent Semantic Indexing (LSI) with 50 SVD components was carried out on the scATAC-seq data. For *cis*-regulatory inference, scglue v. 0.3.2^67^ was used. An additional population of RNA cell barcodes (n = 122) with no association with the experimental design was removed, yielding a total of 14,375 cells for the regulatory inference. Gencode Human Release v.41 was used to annotate the RNA HVFs with promoter regions. The putative connections between the RNA and ATAC HVFs were then modeled as a genomic distance graph, which was constructed to link these HVFs to ATAC peaks within a 150 kb window, with connections weighted by a power-law function to account for genomic contact probability. The ATAC peaks reachable from the RNA HVFs in the guidance graph were marked as HVFs. The HVF graph was then used for model training with the scglue framework on the Nvidia v100 graphics processing unit (GPU), using PCA and LSI embeddings for RNA and ATAC layers, respectively. The learning rate was set to 0.00002 and latent dimensions to 50. Shortly, scglue encodes a common low-dimensional embedding of the data modalities from the graph, which is used for computing cosine similarity of the connections in the graph. This serves as a regulatory score for peak-gene pairs. Empirical *p*-values are computed by comparing regulatory scores to a NULL distribution of randomly shuffled feature embeddings. scglue also generates an integrated latent representation of the data modalities, which was used for generating a UMAP representation of the data.

TF regulons were inferred using the *GRNBoost2* algorithm (arboreto v.0.1.6)^68^ to construct a co-expression network from expressed TFs based on JASPAR2022^69^ motif hits and highly variable genes. The TF-gene pairs were further pruned with pyScenic’s (v.0.12.1)^70^ *ctx* step with rank threshold of 400 to infer the direct targets for each TF, utilizing the *cis*-regulatory ranking of peak–gene connections filtered based on Benjamini-Hochberg *q*-value (< 0.1) to model the TF–peak connections based on the TF motif hits on ATAC peaks and proximal promoters (within +/- 500bp of the TSS) of the genes.

### Inference of TF regulon activity within single cells and computation of metacells

The TF regulons were used as input gene sets for AUCell v.1.24.0 R package^68^, which was to compute the activity score for the regulons. Shortly, AUCell first ranks all genes within each cell by their expression levels, then looks at how many genes from the input set appear high in this ranking, using Area Under the Curve (AUC) score to reflect, how well a gene set is represented among the highly expressed genes (a quantile provided by the user, set to 10 %) in that cell, focusing on the TFs expressed in at least 2 % of the cells with a minimum of five target genes in order to focus on most prominent and robust regulons (n = 23).

To evaluate the activity of the TF regulons within cellular subpopulations, single cells in scRNA-seq data were aggregated into metacells using SEACells v. 0.3.3^4^. Shortly, SEACells uses an archetypal analysis to aggregate single cells into cell states, first applying an adaptive Gaussian kernel to a k-nearest neighbor graph of Euclidean distances between cells to convert the distances into similarities, and then using the kernel matrix for the archetypal analysis using linear decomposition, aiming to summarize the data to summary points based on how similar or close the data points are to each other. The metacell membership is identified based on the maximal values of cells within the archetype in an embedding matrix. Metacells were mapped from the scRNA-seq data to the scATAC-seq data using the *transfer_labels* function in scglue with the combined embedding.

### Exploratory analyses of the TF regulons

The *Enrichr* wrapper of GSEApy v. 1.1.3 was used to analyse the enrichment of the TF regulon target genes with MSigDB Hallmark gene sets^71^ and Reactome^72^ pathways with FDR threshold of 0.05. AUCell was used for computing the gene set activities for single cells, except for the Persist signature, for which *scanpy.tl_score_genes* was used.

To investigate the similarity relationships between the metacells based on the TF regulon activity, we computed a pairwise Euclidean distance matrix using the mean regulon activity scores for each metacell. Distances were calculated using the *pdist* function from Scipy v. 1.11.2. Hierarchical clustering was performed using Ward’s method. The resulting clustering structure was compared with metacell-level cell cycle phase and sample composition. Clusters with higher contribution of S and G2M phase cells were classified as *proliferating* and clusters enriched with G1-phase cells were classified as *non-proliferating*. The non-proliferating cells were retained to analyze the overall pathway activity shifts across the samples. The z-score of DE genes with threshold of *p*_Adj_ < 0.01 and |log2FC| ≥ 0.5 for each dataset were ranked using a jitter-based approach to break ties for rank-based gene set enrichment analysis (GSEA) for MSigDB Hallmark gene sets. The AUCell scores for the gene sets with FDR < 0.05 were used for visualization as heatmap.

Biological relevance of the TF regulons was evaluated using Gene Regulatory Network Performance Analysis (GRaNPA, v. 1.0.4)^26^ on the differentially expressed genes (DEGs) comparing LNCaP-Enza48h, Res-A and Res-B to the parental LNCaP line with *scanpy.tl.rank_genes_groups* (|Log2FC ≥ 0.5|, *p*_Adj_ < 0.05) and the full network of 44 TF regulons. GRaNPA applies a random forest-based regression model to evaluate how well the TF–gene connections are able to predict differential expression driven by TFs in cell type-specific manner. The model is trained to predict gene-specific DE values (here, log2FC) based on the TF–gene connections using 10-fold cross validation. The performance is measured by assessing *R*^2^ of the cross-validation against that of a randomly permuted network. Here, the *R*^2^ values of the real network were 0.19, 0.23 and 0.32 vs. the permuted network 0.06, 0.03 and 0.07, for Enza48h, Res-A and Res-B, respectively.

The TF motif analysis was performed in Signac. Position frequency matrices (PFMs) for the CORE collection were retrieved with the *getMatrixSet()* function from the JASPAR2022 package v.0.99.7, filtering for human motifs and the latest version only. A motif count matrix was generated with *CreateMotifMatrix()* and per-cell motif activity deviation Z-scores were obtained from *RunChromVAR()* function with the hg38 genome.

### Reconstructing *trans*-regulatory programs

To capture broader *trans*-regulatory programs associated with the TF regulons, we first inferred co-expression modules with hdWGCNA (v. 0.4.10)^73^. Genes with mean absolute deviation (MAD) of 0 were ignored. The LogNormalized metacell-to-gene matrix was then used for constructing the coexpression modules with the automatic soft power selection (28 for this dataset). Ignoring genes that were not grouped into any module, we next proceeded to check for the overlap co-expression modules with the TF regulons. The significance of the overlaps was evaluated with Fisher’s exact test.

To translate our findings into gene sets suitable for clinical testing, we defined two non-overlapping networks: an ARS- and stress-linked Enza-sensitive program and a SOX4-linked Enza-resistant program. These networks were constructed by retaining DEGs in datasets with *scanpy.tl.rank_genes_groups* for the regulons and integrating them with the genes from the overlapping co-expression modules. To robustly capture SOX4-linked mechanisms in both single-cell and bulk transcriptomic datasets, we applied a stringent threshold (log2FC > 1, *p*_Adj_ < 0.05) for inclusion of DEGs (Res-A/B vs. LNCaP/LNCaP-Enza48h) in the resistant network. This ensured that only strongly perturbed SOX4 targets were retained, improving robustness for cross-platform validation. Because *SOX4* expression positively correlated with the expression of its target genes (**Fig. 1g**), we included *SOX4* within the SOX4 regulon for all downstream analyses in clinical datasets. For ARS- and stress-associated regulons, we used a broader threshold (log2FC > 0.5, *p*_Adj_ < 0.05) to capture wider transcriptional consequences relevant to ARSI sensitivity (LNCaP-Enza48h vs ResA/B, LNCaP vs. rest). Importantly, we selected this strategy to preserve the network-like logic of the original single-cell data while enabling direct translation to patient cohorts. To assess the behavior of ARS- and stress-linked connections beyond ribosomal biogenesis and canonical androgen response, we excluded ribosomal genes and MSigDB Hallmark Androgen Signaling genes from downstream analyses in clinical datasets.

The ARS- and stress-linked sensitivity program, DE-filtered SOX4 regulon, SOX4-linked resistance program, and additional signatures used for clinical validation, including those associated with cell states (Persist, AR activity, Liu PTEN loss) as well as cell lineages and disease subtypes (Basal, Ductal Luminal, Luminal, NE identity, Beltran NEPC) are reported in **Supplementary Data 2**.

### Additional scRNA-seq data processing

The additional scRNA-seq data derived from patients with treatment-naïve PCa, metastatic hormone sensitive PCa (mHSPC) on ADT and CRPC^14^ was downloaded from GEO (accession number GSE274229) and processed with scanpy according to Lyu *et al*, 2025. Due to most CRPC epithelial cells being from one sample, we focused our analysis on the treatment-naïve PCa and ADT-treated mHSPC cells.

The single-cell multi-condition differential expression analysis was performed with Latent Embedding Multivariate Regression (LEMUR, v. 1.0.5)^74^. The LEMUR model uses matrix factorization to learn condition-specific linear subspaces while integrating cells from different conditions into a shared latent space. A regression component links the known covariates (condition, batch, encoded in a design matrix) to subspaces by defining the condition-specific rotation in the common space. A parametric model is then used to predict the expression of each gene in the cells’ latent coordinates under different conditions. Gene expression change Δ is then computed between conditions from these predictions. We used 15 latent dimensions to predict the effect of disease phenotype in epithelial cells, with *align_harmony()* wrapper (*harmony* v. 1.2.4) applied to improve the between-condition alignment in the shared space. Cell lineage-specific PTEN activity association was evaluated by assigning the epithelial cells into low and high PTEN activity groups based on the median AUCell score of the Liu PTEN loss signature.

The scRNA-seq counts of CRPC needle biopsies^12^ were downloaded from Gene Expression Omnibus (GEO, accession number GSE137829). Raw counts were processed with Scanpy. Quality control of the data was done to keep cells with ≤9000 and ≥ 500 expressed genes and < 10 % of mitochondrial counts, and genes expressed in ≥ 15 cells. *Seurat V3* method^75^ was used to select highly variable genes (n = 2000) for PCA. Data was normalized by total counts over all genes in each cell, logarithmized and scaled, centering each gene to have a mean of 0 and scaled to unit variance.

The Leiden clusters of both datasets were annotated with cell type-specific marker genes (**Supplementary Data 4**). For Dong et al 2020, the single epithelial cells were assigned into high and low SOX4 regulon and SOX4-linked resistance groups based on the z-scored AUCell score (high > 0, low ≤ 0), reflecting relative regulon activity within the epithelial compartment. Epithelial lineage contributions were quantified for each group. For selected epithelial subtypes (club-like and basal cells in the Lyu et al 2025 and luminal-like cells in Dong et al 2020 dataset), the SOX4-linked resistance group was defined again using stricter, quartile-based activity thresholds to better capture the within-population variation. The DE analysis for cells with high vs. low SOX4-linked resistance was carried out, and z-score of DE genes with threshold of *p*_Adj_ < 0.01 and |log2FC| ≥ 0.5 were ranked using a jitter-based approach to break ties and used for the GSEA analysis with GO Biological process and Reactome pathways. The enriched gene sets with FDR *q*-val of < 0.05 and number of leading-edge genes ≥ 5 for Lyu et al 2025 are reported in **Supplementary Data 5**.

### Linear regression analysis

Multivariate linear regression models were used to assess the independent contributions of the regulatory programs and signaling pathways to gene set activity and epithelial lineage. Models were fitted using Ordinary Least Squares (OLS) regression (*statsmodels* v. 0.14.0). Categorical variables were included where appropriate.

To account for the limited sample size, heteroscedasticity-robust standard errors (HC3) were used. Multicollinearity between predictors was assessed using variance inflation factors (VIFs), with all values < 3, indicating no substantial collinearity.

### Bulk RNA-seq data processing

The TPM normalized RNA-seq counts from Enza responder and non-responder patients^31^ were log2(x+1) transformed. Gene set enrichment scores were obtained with the R *GSVA* package (v. 1.50.0). To assess differences in pathway activity between response groups, a gene set-level linear model was fitted using the limma framework (v. 3.58.1). The *p*-values were computed using empirical Bayes moderation of the standard errors was applied to improve the statistical power by stabilizing variance estimates. To examine associations with patient outcomes, samples were assigned to high and low activity groups based on median GSVA score. Overall survival was modeled using Kaplan-Meier curves with log-rank significance testing, implemented via the *survival* package (v. 3.7-0). Figures were generated with *survminer* v.0.5.0.

Processing of additional clinical RNA-seq datasets was conducted in Python. Transcripts per million (TPM)-normalized data from CRPC tumors^3^ was downloaded from GEO (accession number GSE126078) and log2(x+1) transformed. Mutation metadata was obtained from Roudier *et al*, 2025. Additional z-score normalized FPKM (Fragments Per Kilobase of transcript per Million mapped reads) RNA-seq count matrix from CRPC tumors by the SU2C/PCF Dream Team^1^ were downloaded from cBioPortal (https://www.cbioportal.org/study/summary?id=prad_su2c_2015). Log2(x+1) transformed RSEM normalized RNA-seq count data from primary prostate tumors by TCGA^32^ was downloaded from the University of California, Santa Cruz UCSC Xena Browser, along with the CNV data estimated with GISTIC2 method (https://xenabrowser.net/datapages/?cohort=TCGA%20Prostate%20Cancer%20(PRAD)&removeHub=https%3A%2F%2Fxena.treehouse.gi.ucsc.edu%3A443). Log2(x+1) transformed or z-score normalized FPKM counts were used with *gseapy.gsva* (v. 1.1.). The cell lineage-defining signatures were sourced from Hu *et al*, 2025a, except for the NE-like signature^76^, which had undergone further filtering of overlaps between other cell lineage-defining signatures^8^.

To evaluate associations between SOX4 activity and epithelial lineages, the SU2C tumors were assigned to low and high groups for SOX4 regulon and SOX4-linked resistance based on the z-scaled GSVA score (high > 0, low ≤ 0). The dominant cell lineage for the tumors was likewise assigned based on the highest z-scaled GSVA score among epithelial lineage-specific signatures. Distribution of the dominant lineages within the high and low SOX4 activity groups was assessed using the *chi*-square test, followed by lineage-specific Fisher’s exact tests.

### TF motif and footprinting analyses

Differential motif activity between resistant and parental cells was performed on ChromVAR deviation z-scores with *FindMarkers()* in Signac using Wilcoxon Rank Sum test (Mann-Whitney U test) with the average motif activity difference between groups.

For TF footprinting analysis we applied scPRINTER framework v. 1.2.1^77^ on Python 3.11. Peak calling for model training was done using *seq2PRINT* preset, where macs2 summit calls are first padded to 128 bp windows and used for iterative peak merging. After resolving the overlapping peaks, the final peak set is expanded to 1000 bp width from the summits. For the explorative motif analyses, preset *ChromVAR* was used, using the same peak set generation strategy, but the iterative peak filtering is performed at 800 bp resolution, after which the final peaks are resized to 300 bp width. Peaks were called for each sample separately, with the iterative peak merging done across samples with FDR threshold of 0.01.

The seq2PRINT model was first trained on the Nvidia A100 GPU to learn general TF footprint patterns of the samples on bulk level. For each metacell, the model was then fine-tuned employing the low-rank adaptation of large models (LoRA) with 5-fold cross validation, using PCA-reduced cell embeddings derived from ChromVAR deviation z-score computed with scPrinter’s GPU implementation. Background peaks were sampled for each peak using the nearest neighbor method (‘nndescent’) with 250 neighbors per peak. Motif matches were then scanned across the peaks using the human TF motifs from functional inference of gene regulation (FigR) framework^78^, with *p*-value threshold of 5 x 10^-5^.

TF binding prediction was carried out for the SOX4 regulatory region based on the sequence attribution scores from seq2PRINT together with JASPAR2022 core motif matches. Sequence attribution score is the base-to-base measure of importance for the prediction of the model. The regulatory region for SOX4 was selected based on the same criteria as for *cis*-regulatory inference, by considering the genomic window within 150 kb around the gene. TF binding score (denoting the binding probability of each TF) for peak sequences overlapping these genomic windows was computed on the level of metacells, focusing only on the TFs expressed in the scRNA-seq layer. Additionally, sequence attribution scores from the accessibility count layer were used for *de novo* motif calling, using modisco-lite v. 2.4.0^79^ wrapper in scPRINER to identify regions with high scores (‘seqlets’), which are clustered into negative or positive *de novo* motif groups based on their similarity. These *de novo* motifs are then matched with known motifs using *tomtom* from memelite v.0.2.0^80^, after which the de novo motif occurrences are called with finemo v. 0.30^81^. Motif matching analysis for the scPrinter peaks was done using the HOCOMOCO v.14 TF motif collection^82^. Motif scores and motif matches across the peaks within the SOX4 regulatory region were computed with motifmatchr R package (v. 1.24.0). The genome track shown in **Fig. 4f** was generated using *pyGenomeTracks* (v. 3.9). The annotations for AR motif matches and the scPRINTER *de novo* motif patterns and known motif binding events were used to assess potential motif co-occurrence by merging the motif intervals within 100bp windows using pyranges (v. 0.1.4). Gene annotations were sourced from Gencode Human Release v.41. The scATAC-seq and scRNA-seq BAM files were converted into CPM-normalized bigWig tracks with *bamCoverage* function from deepTools (v. 3.5.5).

### Statistics and reproducibility

Statistical testing and multiple testing corrections were performed with *scipy.stats* v. 1.11.2 and *statsmodels.stats.multitest* v. 0.14.0, respectively. Unless stated otherwise, all tests were two-sided and *p*-values were adjusted using Benjamini-Hochberg procedure. Overlapping genes were removed when calculating scores for correlation analyses between gene signatures and for multivariate linear regression analyses between predictor and response gene sets.

## Data availability

The scRNA-seq and scATAC-seq data used for the regulatory inference is available in the Gene Expression Omnibus (GEO) archive under accessions GSE168669. Other publicly available scRNA- seq data were from patients with treatment-naïve PCa, metastatic hormone sensitive PCa (mHSPC) on ADT and CRPC^14^ (accession number GSE274229) and from six CRPC needle biopsies^12^ (GSE137829), as well as bulk RNA-seq data from the Labrecque CRPC cohort^3^ (GSE126078), with mutation metadata obtained from Roudier *et al*, 2025, CRPC cohort by the SU2C/PCF Dream Team^1^ (https://www.cbioportal.org/study/summary?id=prad_su2c_2015) and from primary prostate tumors by TCGA^32^, along with the CNV data (https://xenabrowser.net/datapages/?cohort=TCGA%20Prostate%20Cancer%20(PRAD)&removeHub=https%3A%2F%2Fxena.treehouse.gi.ucsc.edu%3A443).

## Code availability

Code to reproduce the results is available at https://github.com/sinihak/PCa_Enza_res_scGRN.

## Acknowledgements

The authors wish to acknowledge Tampere Center for Scientific Computing (TCSC), Finland, and CSC–IT Center for Science, Finland, for computational resources. We want to thank Päivi Martikainen, Hanna Selin, and Karishma Sajnani for assistance in experiments. S.H. has received funding from Finnish Cultural Foundation and University of Tampere Foundation sr. M.M. acknowledges the Research council of Finland (347543), Sigrid Jusélius foundation, and Instrumentarium Science foundation. A.U. wishes to thank the Research Council of Finland project no. 349314; Cancer Foundation Finland; Norwegian Cancer Society project no. 273672 –2023 and Tampere Institute for Advanced Study. M.N has been funded by the Academy of Finland Center of Excellence program (project no. 312043), the Sigrid Jusélius Foundation, and the Cancer Foundation Finland.

## Author contributions

S.H. conceived and performed the computational analysis and wrote the paper. A.P. performed experiments. A.K. assisted in computational analysis. A.U. assisted in data interpretation and writing of the paper. M.N. and M.M. supervised and directed the project, assisted in writing of the paper, and secured financial support.

