## Supplementary Figures 1-5 for "AR suppression rewires a SOX4 regulatory program to promote enzalutamide resistance in prostate cancer"

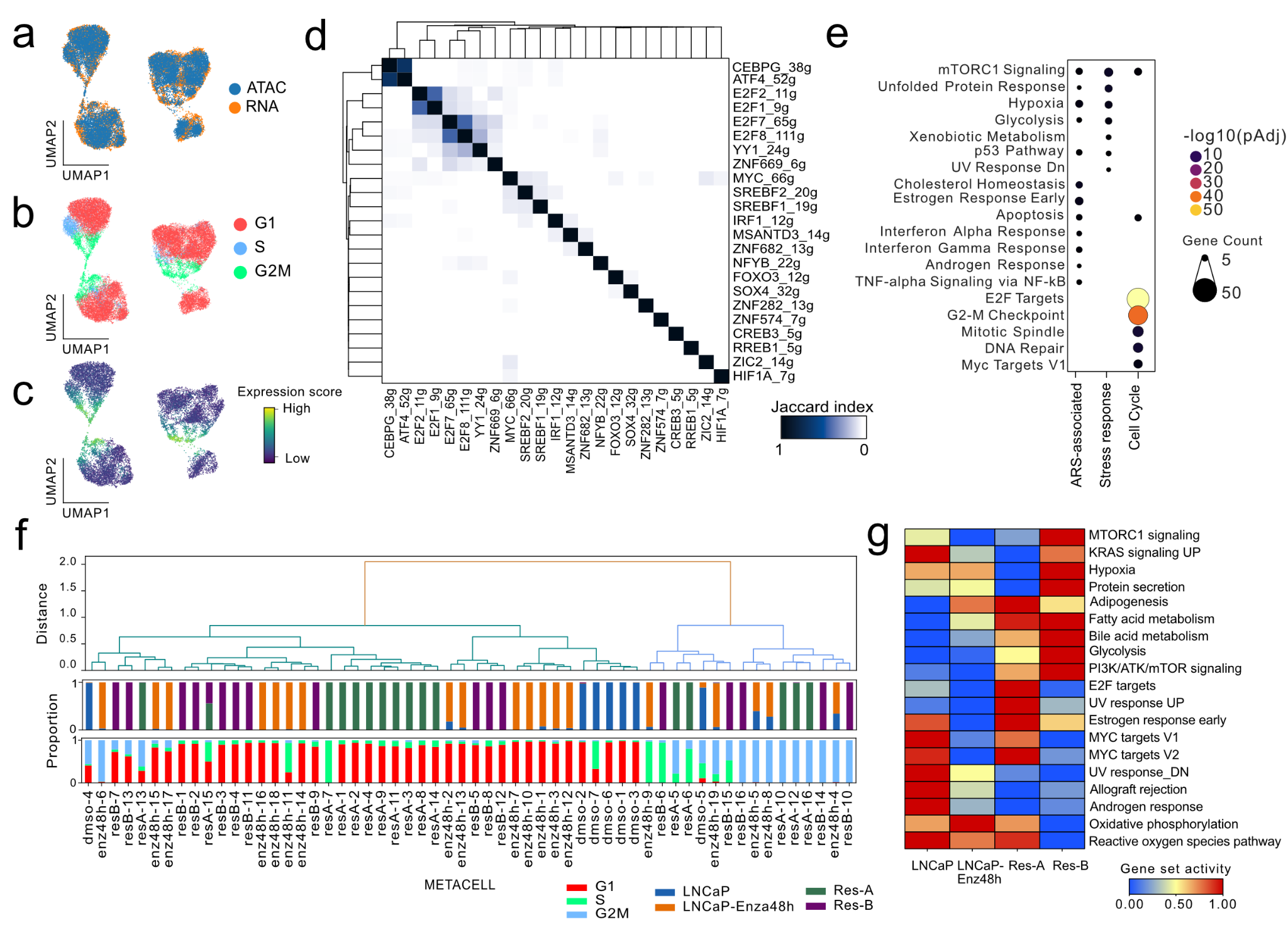

**Supplementary Figure 1.** **a–c** UMAP colored by the **a** integrated data layers **b** cell cycle phase and **c** expression score of the Persist signature in the RNA-layer **d** Jaccard index heatmap of the gene-level overlap of the TF regulons. **e** MSigDB hallmark gene sets the TF regulon groups ARS-associated, Stress Response and Cell Cycle regulons are enriched in (permutation test p_Adj_ < 0.05). The Resistant regulons were not significantly enriched in MSigDB hallmark gene sets and are therefore not shown. **f** Hierarchical clustering of the metacells based on the TF regulon activity scores using Ward’s method. Bar plot annotations below the dendrogram indicate the proportion of single cells within each metacell assigned to different cell cycle phases (G1, S, G2M) and samples. Clusters enriched for S/G2M phase cells were classified as *proliferating* (light blue), whereas G1-enriched clusters were classified as *non-proliferating* (green). **g** Activity heatmap of the significantly altered MSigDB hallmark pathways identified by gene set enrichment analysis (GSEA; FDR *q*-val < 0.05) across datasets, considering only cells within *non-proliferating* metacells.

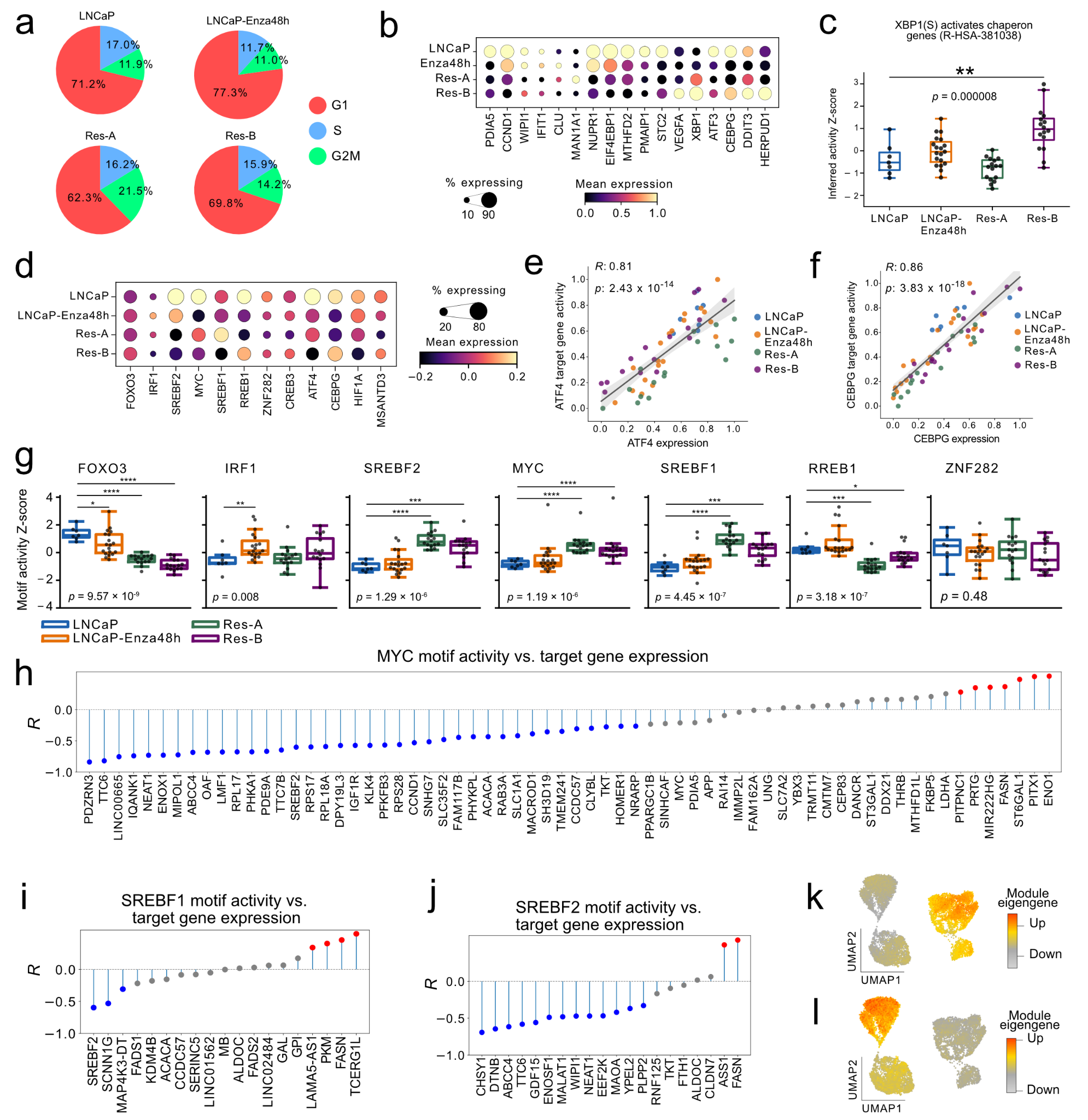

**Supplementary Figure 2. a** Pie charts showing the percentage of cells in each cell cycle phase (G1, S, G2M) across samples in the *in vitro* model. **b** Scaled gene expression of unfolded protein response-associated genes in the ARS-associated and Stress Response regulons. **c** Inferred activity of Reactome Pathway Database XBP1(S) activates chaperone genes (R-HSA-381038) gene set on metacell level. Metacells were grouped based on the dominant sample within each metacell. Value represents z-score normalized activity score. Kruskal-Wallis test *p*-value is displayed, with the asterisk indicating Mann-Whitney U test *p*-value ** < 0.01. **d** Expression z-score of stress response and ARS-associated regulon TFs. **e–f** Correlation scatter of **e** ATF4 and **f** CEBPG expression and corresponding regulon activity across metacells**,** colored by the dominant sample within each metacell. *R* represents the Spearman correlation coefficient and *p* the associated Student's t-test *p*-value. **g**. Motif activity of the ARS-associated regulons in metacells. Vvalue represents the z-score normalized activity score. Kruskal-Wallis *p*_Adj_ is displayed ( ****p*_adj_ < 0.001, *****p*_adj_ < 0.0001, Mann-Whitney U test). **h–j** Lollipop plot of Spearman correlation coefficient between the expression of **h** *MYC* **i** *SREBF1* and **j** *SREBF2* motif activity and their genes. Grey dots denote correlations with *p* ≥ 0.05. The **k–l** Co-expression module **k** sc-M1and **l** sc-M9 eigengenes (the first principal component of the module gene expression) represented as UMAP.

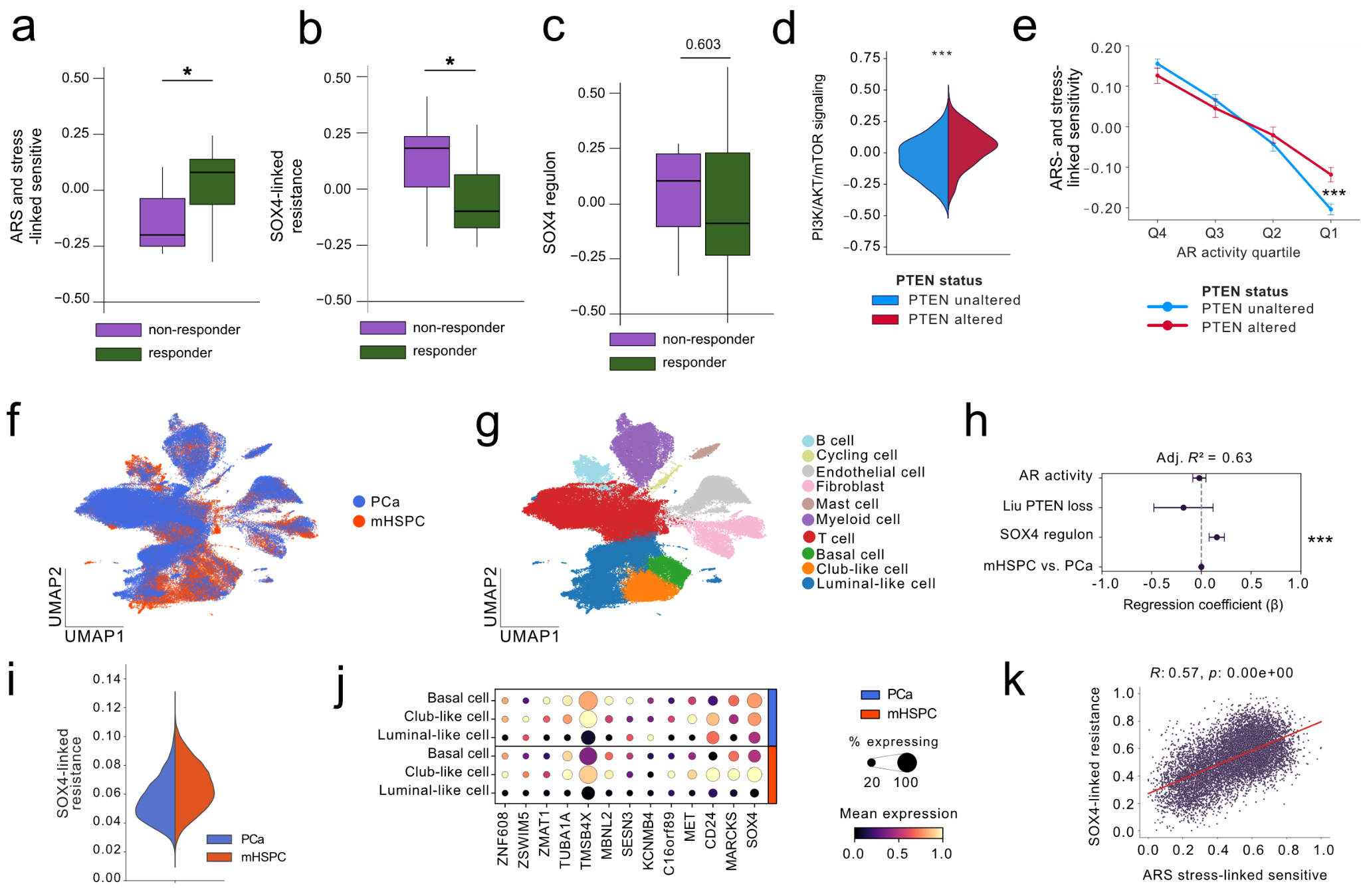

**Supplementary Figure 3.** **a–c** GSVA score of **a** ARS- and stress linked sensitivity **b** SOX4-linked resistance and **c** SOX4 regulon in Enza-treated CRPC patients (n = 25) classified as responsive or non-responsive based on PSA-levels after 12 weeks of treatment^1^. The asterisk represents *p*-value between groups from the limma moderated *t*-tests with empirical Bayes variance shrinkage (* for *p*-value < 0.05). **d** Violin plot of GSVA scores of PI3K/AKT/mTOR signaling activity in TCGA PCa cohort (n = 491)^2^ stratified by PTEN CNA status. **e** ARS- and stress-linked sensitivity in TCGA PCa cohort across AR activity quartiles stratified by PTEN CNA status. **f–k** Analysis of scRNA-seq data from PCa and ADT-treated mHSPC patients^3^. **f–g** UMAP embedding of colored by **f** tumor type **g** cell type. **h** Ordinary least squares (OLS) regression coefficients for models predicting activity of SOX-linked resistance using SOX4 regulon, Liu PTEN loss signature^4^ activity, AR activity and tumor type (PCa vs. HSPC). Dots show coefficients and error bars the 95 % coefficient intervals (HC3 robust SE). Scores were pseudo-bulked per sample (n = 37), and Adjusted *R*^2^ is shown. **i** Violin plot of **the** SOX4-linked resistance activity in the epithelial cells from PCa and mHSPC patients. **j** Expression of SOX4 regulon in epithelial sub-lineages. **k** Correlation scatter plot of min-max scaled activity of ARS- and stress-linked sensitive connections versus SOX4-linked resistant connections in basal and club-like epithelial mHSPC cells. *R* denotes the Spearman correlation coefficient and *p* the associated Student's *t*-test *p*-value.

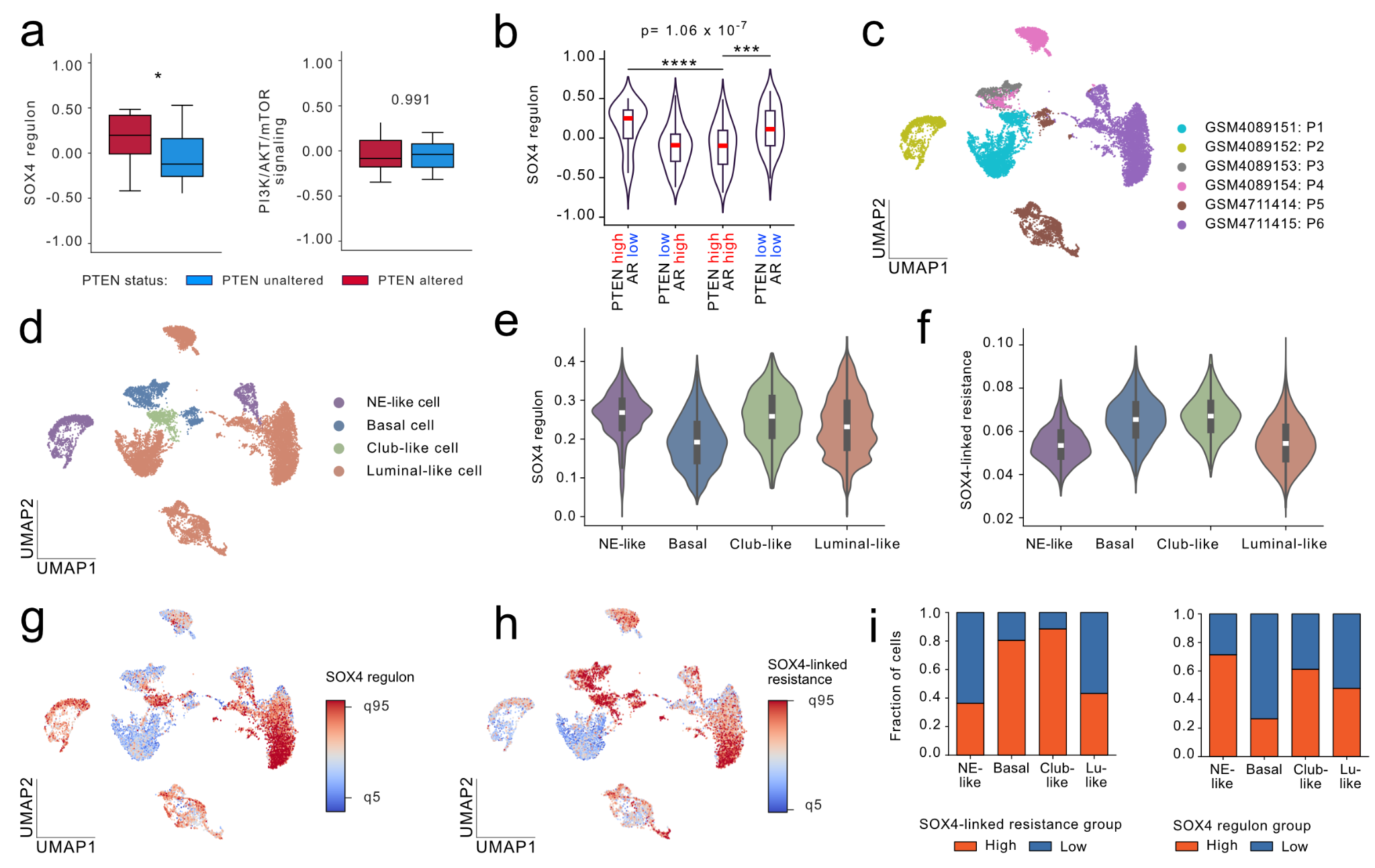

**Supplementary Figure 4. a** GSVA score of SOX4 regulon and PI3K/AKT/mTOR signaling activity in Labrecque 2019 cohort^5^ represented as boxplots, stratified by PTEN copy number status: PTEN altered (n = 26), and PTEN unaltered (n = 20). The asterisk represents Mann-Whitney U test *p*-value * < 0.05. **b** GSVA score of SOX4-linked resistance, and ARS- and stress-linked sensitivity in SU2C CRPC cohort (n = 249)^6^ stratified by AR and PTEN activity status. Groups were defined based on the z-scored GSVA scores (AR low/high corresponded to Z ≤ 0/> 0. As the PTEN signature reflects PTEN loss, higher scores indicate reduced PTEN activity, thus Z ≤ 0/> 0 corresponds to PTEN high and PTEN low, respectively. The Kruskal-Wallis *p*_adj_ is displayed (*** *p*_adj_ < 0.001, **** *p*_adj_ < 0.0001, Mann-Whitney U-test). **c-i** Analysis of CRPC scRNA-seq dataset from prostate tumors^7^. **c–d** UMAP visualization colored by **c** patient and **d** cell type. **e–f** Violin plots of **e** SOX4 regulon and **f** SOX4-linked resistance activity in CRPC cell types. **g–h** UMAP visualization colored by **g** SOX4 regulon and **i** SOX4-linked resistance activity. **i** Stacked barplots describing the cell sub-lineage fraction after stratifying single cells into High and Low SOX4 regulon or SOX4-linked resistance activity groups based on 1st and 4th activity quartiles. Bar height represents the proportional distribution of cell sub-lineage within each activity group

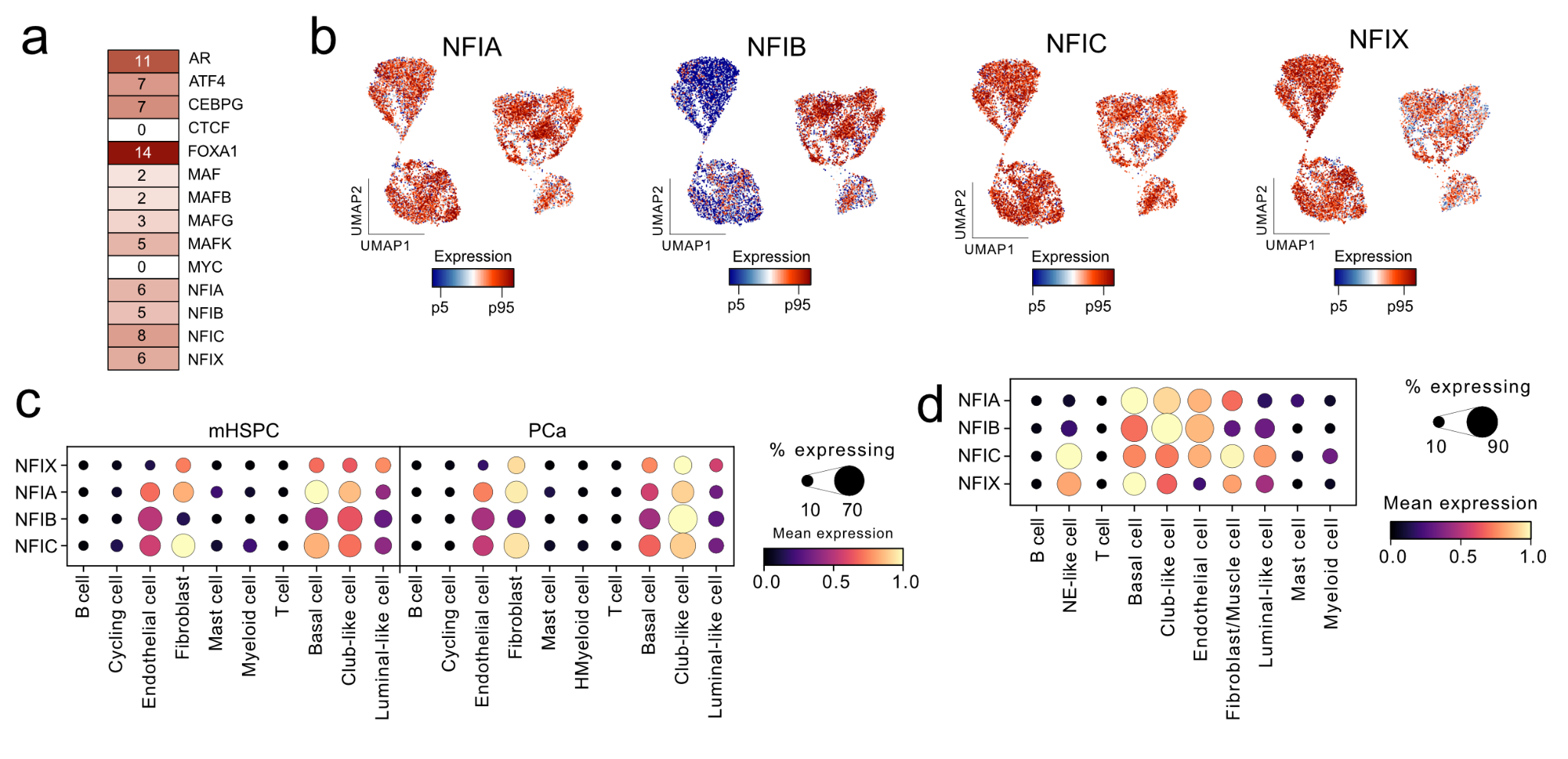

**Supplementary Figure 5. a**. Motif matches for selected TFs at the scATAC-seq peaks overlapping the *SOX4* regulatory region. The analysis was restricted to NFI and MAF family factors, FOXA1, AR, CTCF, ATF4, CEPBG and MYC. **b** UMAPs colored by the expression of NFI family TFs. **c** NFI factor gene expression in scRNA-seq data from mHSPC and PCa patients^3^. **d** NFI factor gene expression in scRNA-seq data from CRPC patients^7^.
